# Universality in the Mechanical Behavior of Vertex Models for Biological Tissues

**DOI:** 10.1101/2022.06.01.494406

**Authors:** Ojan Khatib Damavandi, Sadjad Arzash, Elizabeth Lawson-Keister, M. Lisa Manning

## Abstract

Simple vertex models, where the cell shape is defined as a network of edges and vertices, have made useful predictions about the collective behavior of confluent biological tissues, including rigidity transitions. Quite a few different versions of vertex models have appeared in the literature, and they propose substantial differences in how the mechanical energy depends on vertex positions, yet all of them seem to make correct predictions. To understand how this is possible, we search for universality in the emergent mechanical behavior – including the shear modulus defined in the limit of zero strain rate and the viscoelastic response at finite strain rates – of six different vertex models. We identify a class of models with a well-defined shear modulus, and demonstrate that these models all exhibit a cross-over from a soft or floppy regime to a stiff regime. While the parameter that controls the crossover is different in each model, we find that the observed cell shape index (the ratio of the cell perimeter to the square root of the cell area) is a good observable order parameter for the crossover. We also find that the finite strain-rate viscoelastic response of all models exhibits a universal scaling with frequency, following the Zener model in the rigid phase and Burgers model in the fluid phase. This suggests there is a broad class of vertex models with universal mechanical features, and helps to explain why many different vertex models are able to robustly predict these features in experiments.

## I. INTRODUCTION

In biological processes ranging from embryonic development to cancer metastasis, recent work has emphasized that the collective behavior of cells plays an important role [1–3]. Many of these animal tissues undergo a fluid-solid transition which is important for their function [4–6]; tissues fluidize in order to allow large-scale deformation and then solidify to maintain functional shapes and support shear stresses.

Theoretical and computational work in concert with experimental observations have identified several mechanisms [7] that can generate fluid-solid transitions tissues, including an increase in cell density to increase the number of contacts and generate percolating rigid cluster [4, 8], a tension-driven rigidity transition in the vertex model [9, 10], change in active fluctuations [11].

Quantifying the onset fluid-solid transitions in disordered materials is a non-trivial problem, so these works have also had to develop heuristic tools to do so. The standard method is the shear modulus, which is the linear response of the system to an applied shear strain in the limit of infinitely slow driving, a definition that works best at zero temperature when there are no competing timescales [12]. In glassy systems at finite temperature, there can be quite large competing timescales, and so in practice experiments on glassy materials define a material to be solid when its measured viscosity exceeds a specified threshold [13]. Similar methods in simulations of active glassy matter place a threshold on the measured diffusion constant [14] or self-overlap function [15].

Given these competing timescales, a more robust and quantitative approach is to study the full response of the tissue as a function of the rate of deformation [16]. Many biological tissues exhibit a viscoelastic response that changes as a function of the frequency of an oscillating deformation. These responses have been measured in both experiments [4, 17] and in simulations of the standard vertex model [18, 19] and active vertex model [3]. These works find that tissues exhibit a complex viscoelastic rheology.

Comparison of both the zero-strain-rate and finite strain-rate responses predicted by coarse-grained models with experimental observations demonstrate that the models are quite successful in capturing fluid-to-solid transitions in tissues. Particle-based models have made accurate quantitative predictions for the relationship between cell density and network connectivity and tissue fluidity in the developing zebrafish embryo [4, 8], and vertex models have made accurate quantitative predictions for the relationship between cell shape and tissue fluidity/remodeling in human bronchial epithelial tissues [5], fruit fly embryos [6] and MDCK monolayers [20], as well as the full viscoelastic response in the zebrafish tailbud [17, 19].

In this work, we focus on vertex models, which assume that a confluent epithelial monolayer with no gaps or overlaps between cells can be described as a network of edges and vertices [9, 21, 22]. In a standard version of the model, the energy functional that describes the tissue is written as a sum of two terms: a quadratic penalty on the cross-sectional area and cross-sectional perimeter when those quantities deviate from their target values. These terms naturally arise when considering volume incompressibility, cortical tension and adhesion, and active contractility in epithelial monolayers [9].

Much of the work on vertex models has focused on 2D systems, as epithelial monolayers occur regularly in development and disease processes and the experimental data is fairly straightforward to compare to models. Although some work has suggested that similar rigidity transitions occur in 3D tissues as well [4, 23, 24], for simplicity and to make clear connections with previous literature, we also focus here on 2D models.

While the tension-driven rigidity transition appears to be a quantitatively predictive theory for several experimental systems (mainly in epithelia), the simple vertex model that has been used to study features of the rigidity transition is in some ways clearly too simple. It does not include many features that are relevant for real biological tissues. Therefore, an open question remains: **why is it predictive at all?** One possible answer to this question is that there is a larger set of models that all exhibit an underlying rigidity transition with universal features. To understand whether something like this is possible, it is useful to examine variations of vertex models that have been proposed in the literature.

There are different classes of variations to vertex models. A class of relatively mild variations involves altering the initial conditions or the dynamics of the vertex model to add specific biologically relevant details, and studying the resulting rigidity transition. One study investigated the effect of rosette formation [25], finding that an increasing number of rosettes further rigidifies a tissue. A related series of studies investigated vertex models with so-called “T1 delay times”, where it is assumed that molecular-scale processes could impede individual cell-rearrangements over a characteristic timescale [26, 27]. In those models, delay times could enhance the rigidity of tissues and also generate cellular streaming patterns consistent with experimental observations. Another set of studies investigated how heterogeneities in cell stiffnesses affect vertex models, finding that rigidity percolation of a sub-population of stiff cells could be used to explain the onset of rigidity in that case [28].

A class of more drastic variations involves fundamentally altering the vertex energy functional itself by adding different or dynamical degrees of freedom. One such work is the active foam model developed by Kim et al [29], which removes the second-order term that penalizes deviations from a target perimeter. Instead, there is only a linear penalty for variations to the perimeter, similar to the role surface tension plays in foams. Another variation replaces the second-order term for the perimeter with a second-order term penalizing deviations of each individual edge length from a target value, generating a system that is reminiscent of a spring network, albeit still with constraints on cell areas. An even more realistic variation is to allow the edges to have additional degrees of freedom and dynamics that are described by additional differential equations. In work by Staddon and collaborators [30], edges are mechanosensitive springs that possess rest-lengths that can change depending on the tension and strain across the edge. In work by Noll et al [3], edges are springs with rest-lengths that are governed by myosin dynamics, which is, in turn, mechanosensitive and governed by tension on the edges.

An obvious question, then, is how this more drastic (and realistic) set of perturbations to vertex models affects tissue viscoelasticity and fluid-solid transitions. Is there still a rigidity transition in such systems? If so, does the transition exhibit similar features to the one seen in standard vertex models? On either side of the transition, does the full viscoelastic response display similar scaling with frequency? If there are universal features in a broad class of vertex models, this could help explain why such simple models are quantitatively predictive in experiments. Non-universal features could help point the way to experiments that can distinguish between models.

Here, we analyze the onset of rigidity and finite-frequency viscoelastic response in a set of six vertex models: the standard vertex model and five others where the energy functional is fundamentally altered.

## II. METHODS AND MODELS

### A. Quantifying mechanical response

#### 1. Shear modulus in the limit of vanishing strain rate

We first develop a framework for comparing different vertex models to one another. The change in free energy of a vertex model can be written generally as

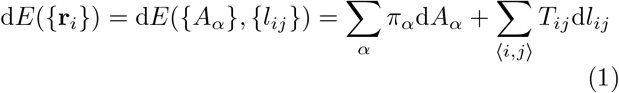

where {**r**_*i*_} is the set of vertex coordinates, {*A*_*α*_} is the area of cell *α*, {*l*_*ij*_} is the length of the edge between vertices *i* and *j*, and 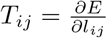 is the tension on the edge between vertices *i* and *j*. Pressure, *π*_*α*_, generally depends linearly on how far cell areas are from their preferred area *A*_0_. The choice of the form of edge tensions can lead to different vertex models with potentially different properties.

As this work focuses on rigidity transitions in vertex models in the limit of zero fluctuations, we use the shear modulus as the metric for rigidity in the limit of zero strain rate. The shear modulus is zero in a floppy or fluid-like system and non-zero in a rigid system, except under special circumstances [31]. The shear modulus *G* is defined as the second derivative of the energy *E* with respect to applied shear strain *γ*, or equivalently the first derivative of the shear stress *σ* with respect to *γ*:

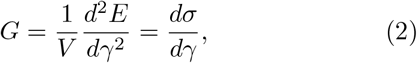

where *V* is the total volume (area) of the system. In practice, we compute *G* in our simulations using the Hessian matrix of second derivatives of the energy with respect to vertex displacements [23]:

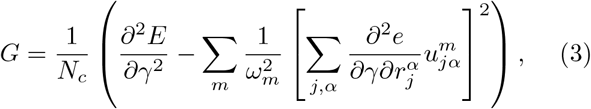

where 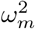 are the non zero eigenvalues of the Hessian *D*_*jα,kβ*_, and 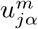 are the corresponding normalized eigen-vectors:

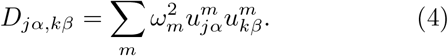

#### 2. Computing linear viscoelastic response at finite strain rate

We are also interested in the mechanical viscoelastic response of vertex models under finite rates of deformation, where there are often rearrangements of the network structure. Viscoelasticity is typically measured by analyzing the response of material to a small amplitude oscillatory strain. Here we briefly review the standard methods we use for computing this response. As with the shear modulus, we focus on the simple shear deformation tensor,

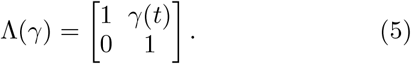

The strain is varied periodically with a sinusoidal function at an angular frequency *ω* as

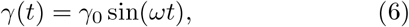

where *γ*_0_ is the maximum strain amplitude. If the viscoelastic behavior is *linear*, the shear stress also oscillates sinusoidally but will be out of phase with the strain [16],

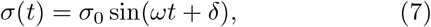

where *δ* is the phase shift. For an ideal elastic solid the phase shift is 0°, and for a Newtonian fluid this phase shift is 90°. Systems with 0° < *δ* < 90° are viscoelastic and the stress response can be written in terms of elastic (or in-phase) and viscous (or out-of-phase) contributions

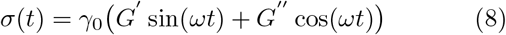

where *G*′ = (*σ*_0_*/γ*_0_) cos(*δ*), *G*″ = (*σ*_0_*/γ*_0_) sin(*δ*) are the shear storage and loss moduli, respectively [32], that are only well-defined when the response is linear [33].

In practice, we compute the storage and loss moduli from the complex modulus *G*^*^ = *G*′ + *iG*″ [32], which is the ratio of the Fourier-transformed stress over strain

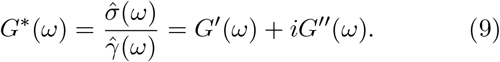

To perform these dynamic oscillations in our vertex models, we use the overdamped equations of motion for vertices, i.e., we neglect inertial effects as is standard for tissue simulations. In this case, the interaction forces will be balanced by the frictional forces from the substrate

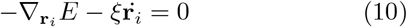

where the first term is the force due to interactions and the second term is a mean-field frictional force with a friction coefficient *ξ*.Other choices for the functional form of the damping term are possible, e.g., proportional to the difference in velocities between neighboring cells, and these would be interesting to investigate in future work. We fix *ξ* in our simulations, which sets the unit of time to *τ* = *ξ/*(K_*A*_*A*_0_). In this work, we report the frequency in inverse natural units, 1*/τ* = (K_*A*_*A*_0_)/*ξ*. In past work by some of us, we performed rheological simulations in the standard vertex model via a deforming droplet, and compared directly to droplet experiments in zebrafish by Serwane et al. [17]. This allowed us to derive an estimate for *τ*, which was about 1 minute [19]. Therefore, a rough estimate for the frequency *ω* = 1 in our simulations is about 1 Hz.

We apply oscillatory simple shear deformations to the vertex models according to Eq. (6). After the affine shear strain in each time step, we evolve the system using the overdamped equation of motion Eq. (10) (See Supplemental Movie [34]). To ensure we probe the linear response, we use a small strain amplitude *γ*_0_ = 10^−7^. We conduct 20 oscillation cycles for each frequency *ω*, discarding the initial 5 cycles to allow the system to stabilize and reach a steady state. We make sure that the system reaches a steady state by analyzing the Lissajous–Bowditch curves, as detailed in the Appendix. Additionally, we utilize a time step that guarantees a minimum of 32 data points within each cycle. We compute the storage and loss moduli using Eq 9, as discussed in more detail in the Appendix.

#### 3. Computing nonlinear viscoelastic response at finite strain rate

Vertex models with dynamical internal degrees of freedom sometimes exhibit significant nonlinearities and transient behavior under the oscillatory tests. This is expected since their state is evolving as we shear them, but it means that we cannot use standard linear response to analyze them. We plot Lissajous–Bowditch curves for such models and find drifting behavior, indicative of a transient state (see Appendix). Thus, to accurately assess the elastic and viscous contributions in dynamical vertex models, we need to introduce an additional parameter known as the *waiting time* (t_*w*_). This parameter represents the duration since the internal dynamics of the models began. At *t*_*w*_, we temporarily halt these dynamics and obtain the system’s minimum energy state to expedite reaching a steady state in the oscillatory simulations. Subsequently, we apply an oscillatory test to this state to compute the storage (G′) and loss (G″) moduli.

Although it is not possible to freeze the internal degrees of freedom in experiments, this approach, often referred to as “rheological aging”, is has been successfully utilized to characterize and understand the viscoelastic response of other nonlinear systems with significant transient responses, including dynamic glassy systems [35, 36] or aging suspensions [37]. It provides a compact and useful description of how the rheology is impacted by transient dynamics. For example, the behavior of these dynamic vertex models in the large *t*_*w*_ limit provides insight into their long-term behavior after extended internal activity.

### B. Models

#### 1. Vertex model

One of the most widely used models, introduced by Farhadifar et al [9] and which we simply call the standard vertex model, assumes that tension on an edge of cell *α* is proportional to *P*_*α*_ − *P*_0_, where *P*_0_ is the cell’s preferred perimeter. For a system that follows this vertex model, the energy integrated from Eq. 1, up to a constant, is

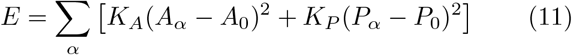

This equation can be non-dimensionalized, identifying two dimensionless parameters, the dimensionless ratio between the area and perimeter moduli *K*_*P*_ */*(K_*A*_*A*_0_), and the cell target shape index 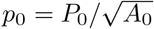.

As most of the previous work to understand the rigidity transition in this class of models has focused on the standard vertex model, we briefly highlight what is known. First, the seminal work by Farhardifar et al [9] demonstrated that there were two ground states of this model – a rigid ordered network and a soft disordered network. Bi et al [10] studied disordered metastable states of the network and found that energy barriers to cellular rearrangements, quantified by statistics of shrinking edges, disappeared above a critical value of the cell target shape index 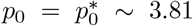. This suggests that the standard vertex model exhibits a floppy-to-rigid transition as *p*_0_ is decreased below a critical value. Quantifying the shear modulus near the transition is quite difficult due to the nearly flat energy landscape, but some of us demonstrated that it, too, disappears in disordered systems for 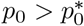 [23].

Interestingly, in ordered ground states of the model, there is a distinction between the onset of rigidity determined by the shear modulus (which occurs at the shape index of a regular hexagon, *p*_0_ ∼ 3.72), and when energy barriers disappear (at 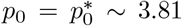). This suggests that there must be non-analytic cusps in the potential energy landscape [38, 39] and that there may be significant differences between the linear (zero strain rate, infinitesimal strain) and nonlinear (finite strain rate, finite strain) rheology of vertex models, which have recently been studied in 2D [39–41] and 3D [42].

Recent studies have also focused on understanding the underlying mechanism that drives the vertex model rigidity transition, as it does not follow a simple Maxwell constraint counting argument [38, 43]. A few studies have demonstrated that a geometric incompatibility between the lengthscale imposed by the target shape index and the cell area (or equivalently, the shape imposed by the boundary conditions) drives the transition [44, 45]. Most recently, we and collaborators have demonstrated that second-order rigidity (penalties that arise only to second order in the constrains) and not first-order rigidity (another name for Maxwell constraint counting) drives the rigidity transition in vertex models[31, 46].

#### 2. Confluent active foam model

As highlighted in the introduction, recent work by Kim et al [29] simulated deformable, adhesive particles and demonstrated that there was an interesting crossover from a particle-like jammed system with holes between cells to an epithelial-like rigid material where cells tiled all space. Importantly, this work used a different energy functional than the vertex model discussed above. Called an “active foam” model, the authors did not include a restoring force on the perimeter (i.e. a term proportional to *P*^2^), and instead assumed that the tension on the edges is simply a constant for all edges, *T*_*ij*_ = Λ. In this work we study the confluent limit of the active foam model, where the density is sufficiently high to ensure that the cells tile all space with no gaps.

This constant tension leads to a simple form of the energy functional:

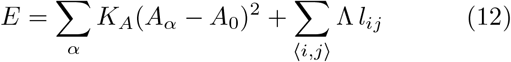

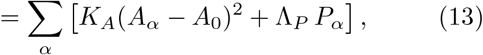

where Λ_*P*_ = Λ*/*2.

Previous work by Farhardifar et al and Staple et al [9, 47] studied this vertex energy for ordered states in a confluent model in the limit that Λ is greater than zero and demonstrated that the system is always rigid. Below, we study the stability of the model when the line tension Λ is zero or negative.

We note that other works explore related changes to the second-order perimeter term of the energy functional. For example, work by Latorre et al [48] investigates a vertex-like model where the second-order restoring term for the perimeter depends on the cell area/volume, so that as the cell area increases the perimeter restoring force decreases. The authors show that such a model, which acts like the active foam model in the limit of large cell size, generates an instability that reproduces many features of cell expansion in epithelial domes.

#### 3. Spring Edge Model

In some epithelial systems, it may be that individual cell interfaces behave as if they are elastic, so that there is a restoring force as the length of that edge shrinks. Such a model might be appropriate for plant cells or animal cells with interstitial ECM and is also sometimes chosen for mathematical convenience [49].

We assume that each edge is a spring with rest (preferred) length *l*_0_. Then, assuming Hooke’s law for the tensions *T*_*ij*_ = 2*K*_*l*_(l_*ij*_ − *l*_0_), and integrating Eq. 1 yields

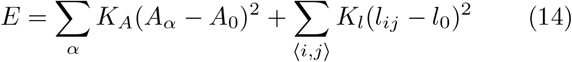

In practice, we simulate the following equivalent expression:

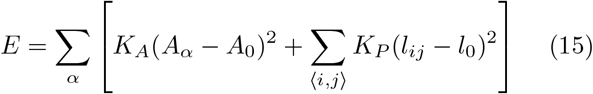

where *K*_*P*_ = *K*_*l*_*/*2 because each edge is counted twice in Eq. 15.

#### 4. Active Spring Model

This model developed by Staddon et al [30] can be thought of as an extension of the spring edge model where spring rest lengths dynamically change to relax the tension. This model is intended to capture active turnover of the actomyosin bundles that give the edges their contractility, replacing strained elements with unstrained ones. It, therefore, posits that the rest length 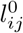 dynamically changes in order to reduce the tension on the edge. It further assumes that only strains above a certain threshold trigger the rest length dynamics, resulting in the following equation of motion:

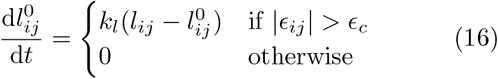

where 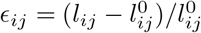.

If *ϵ*_*c*_ = 0, the dynamics will have a completely viscoelastic behavior. This can be seen by taking the time derivative of tension and using Eq. 16: 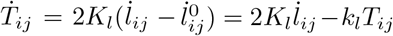 or 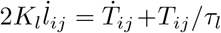, which is simply the Maxwell model of viscoelasticity, with *τ*_*l*_ = 1*/k*_*l*_, the remodeling timescale. The system is expected to fully relax edge tensions at long times. This purely viscoelastic model with *ϵ*_*c*_ = 0 was previously studied in a separate manuscript [50], and the functional form was justified with fits to experimental data during germband extension in the Drosophila embryo. A nonzero *ϵ*_*c*_ means that there can be residual tensions in the network.

We note that Staddon et al. define the tension and energy in terms of strain *ϵ*_*ij*_. However, to be consistent in our definition of tension, we instead define everything in terms of *l*_*ij*_. We expect that qualitatively our results will be similar as the structure of the system at its dynamical steady state, which we discuss below, is the same between these two versions.

#### 5. Tension Remodeling Model

Staddon et al. [30] also define a second “tension remodeling” model that is more consistent with their experimental observations. Specifically, the active tension model is not consistent with the observation that edges can lose memory of their rest length. To generate this feature, the tension remodeling model posits that tension is constant unless strain on an edge reaches a critical value, triggering remodeling of the tension. This can be understood as an extension of the active foam model, where tensions are always constant. They propose the following equation of motion for the tension *T*_*ij*_:

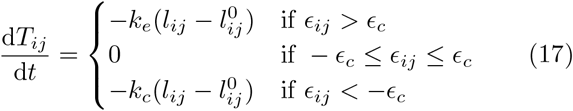

where *k*_*e*_ and *k*_*c*_ are the rates of remodeling under edge extension and compression.

They further postulate that the rest length remodels until the edge is equilibrated:

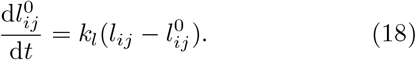

To gain an intuition for this model, first, consider the case where an edge exhibits a strain *ϵ*_*ij*_ < −*ϵ*_*c*_. Then the edge is under compression, and the tension will start to increase (e.g. the edge will become more contractile). Simultaneously, the rest length will shorten and relax the tension until the edge equilibrates at its (shorter) rest length. A similar argument holds in the actively extensile case *ϵ*_*ij*_ > *ϵ*_*c*_.

If instead, the strain is within the nonactive regime for the tension dynamics, even though the rest length continues to change until it matches the actual edge length, the tension remains constant and the force on each vertex is thus fixed, so the vertex positions and edge lengths do not change.

Taken together, this suggests that in all cases the rest length is fixed by the edge length determined by the tension. The energy of this model thus is effectively the foam model, Eq. 13 with Λ → *T*_*ij*_. If we fix the areas, the force on vertex *i* is 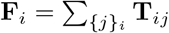.

#### 6. Active Tension Network

A model introduced by Noll et al [3] adds an extra layer of complexity to the tension dynamics by modeling myosin recruitment. The model includes three dynamical processes: 1) vertex dynamics given by energy minimization of Eq. 1, 2) rest length dynamics, and 3) myosin recruitment due to mechanical feedback from the strain at edges.

For their analytics, Noll et al. assume that pressure differences between cells are negligible and thus vertex dynamics are effectively driven by the tension network. With this assumption, the overdamped vertex dynamics (with mobility *µ*) is given by

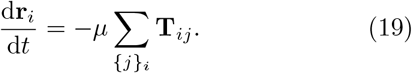

Additionally, it is assumed that each edge acts like an active spring with dynamically changing rest length. Then, 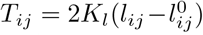 as in the spring-edge model and active spring model, and the energy is given by Eq. 14.

However, unlike the spring dynamics in Eq 16, it is assumed that the fixed point of the differential equation is not at zero strain but rather is achieved when tension on the edge is balanced by the myosin recruited on the edge *m*_*ij*_. In this case, the rest length dynamics is given by

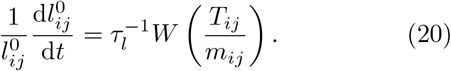

*W*(x) is called the myosin walking kernel, which has the following properties: *W* (1) = 0, *W*′(1) = 1 and the slope of *W* approaches zero near *x* = 0 and for *x* ≫ 1. This means that the fixed point of Eq. 20 is at 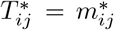. Noll et al. do not consider the *T*_*ij*_ < 0 regime, as they are interested in near-equilibrium behavior where *T*_*ij*_ > 0 due to force balance at each vertex. Linearizing near the steady state gives

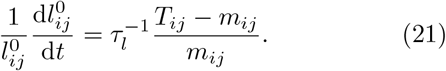

Next, they introduce myosin dynamics. If the myosin field was constant in time, Eq. 21 along with 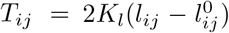 would describe a viscoelastic behavior for the edge length *l*_*ij*_ driven by a shifted (residual) tension *T*_*ij*_−*m*_*ij*_, which would be similar to active spring model with *ϵ*_*c*_ = 0. But since mechanical equilibrium specifies a unique set of tensions, this would mean myosin levels could not be independently prescribed. This is in contrast to experimental observations, as myosin levels have been shown to be dynamic and respond to mechanical cues.

Instead, the authors introduce a simple version of myosin activity that ensures mechanical stability, namely that myosin is involved in a direct mechanical feedback from the edge strain,

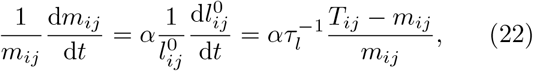

where the second equality holds near equilibrium.

## III. RESULTS

Using the systematic descriptions of six different versions of vertex models developed in the previous section, we proceed to investigate their stability, rigidity, and rheology. Our goal is to understand whether they exhibit a zero-strain-rate rigidity transition and finite-strain-rate rheology similar to those previously described in the standard vertex model.

### A. Standard vertex model

We explore the mechanical response of the standard vertex model under a shear deformation. In the limit of zero strain rate, it is well-established that this model shows a solid to fluid transition as a function of the target shape factor *p*_0_. By computing the zero-frequency shear modulus *G*, researchers have identified the transition point to be in the vicinity of *p*_0_ ≈ 3.9 [10, 44, 51], though the exact critical point may slightly shift due to factors like finite-size effects, the degree of disorder, and the edge threshold for T1 transitions [6, 44].

We compute the finite frequency behavior of such model under oscillatory shear, using methodology described in Sec II A 2 above. This response has already been well-described by Tong et al [18], and we include it here for completeness and comparison to other models.

We analyze both storage and loss moduli as a function of frequency *ω*, and varying *p*_0_ values, capturing the behavior across both solid and fluid phases. The results are shown in Fig. 1.

**FIG. 1.**
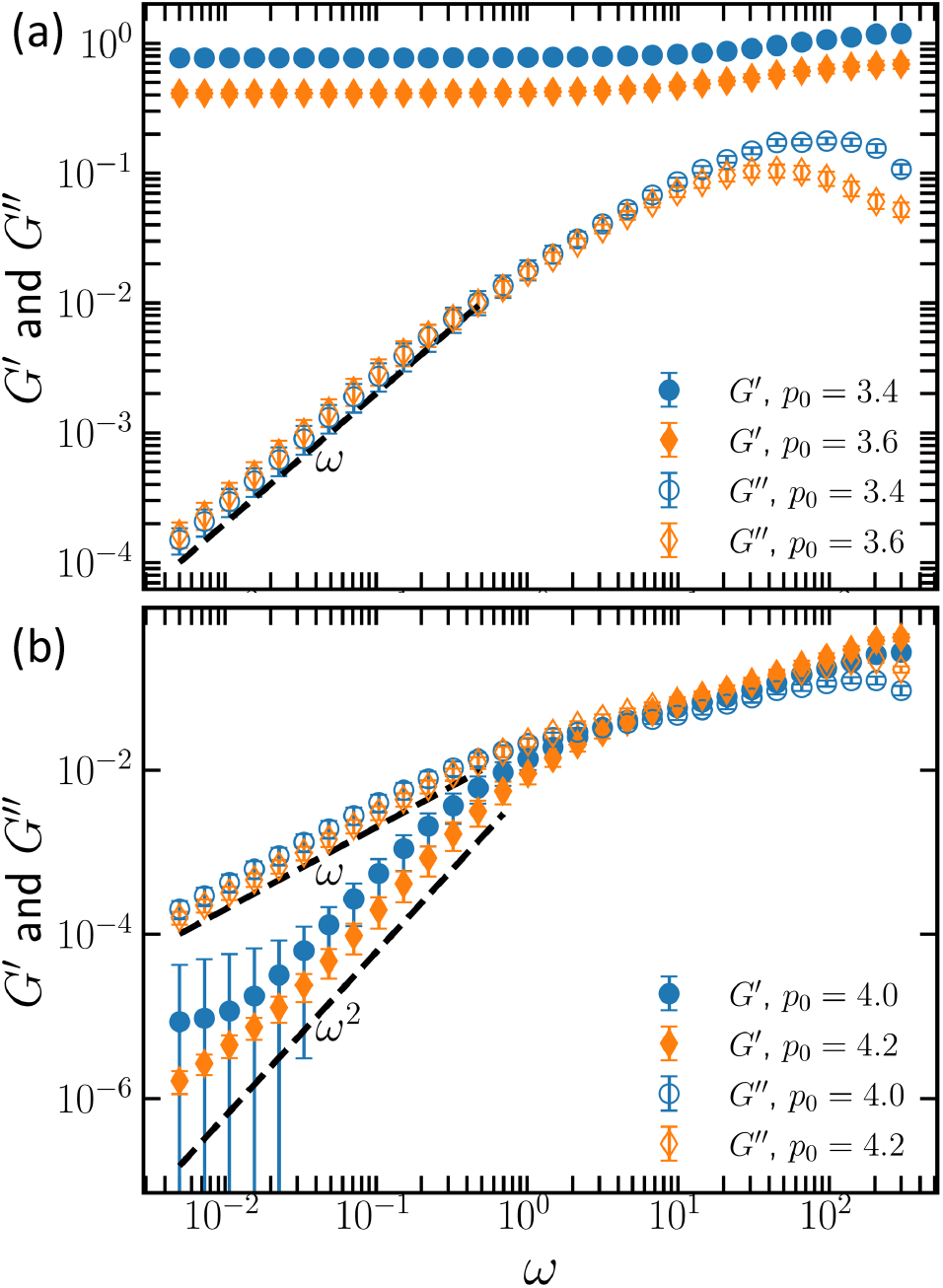
Viscoelastic Behavior of the Standard Vertex Model. (a) The storage and loss moduli, *G*′ and *G*′′, in the solid regime of standard vertex model for two different values of the target shape factors *p*_0_ = 3.4 and 3.6. (b) The behavior of the storage and loss moduli for the standard vertex model in the fluid regime for two different shape factors *p*_0_ = 4.0 and 4.2.

In solid-like phase where the zero-strain-rate modulus *G* is finite and *p*_0_ ≲ 3.9, the rheological response exhibits two distinct regimes, as illustrated in Fig. 1(a). At lower frequencies, *ω*, the storage shear modulus *G*′ exhibits a plateau, while the loss shear modulus *G*″ varies linearly with *ω*. This is consistent with the zero-frequency analysis, since *G*″ vanishes as *ω*→ 0. Conversely, at higher frequencies, *G*′ attains a higher steady value, and *G*″ shows a power-law scaling *G*″∝ *ω*^−1^. This behavior matches the rheology of the Standard Linear Solid or Zener model, a spring-dashpot model to mimic viscoelasticity of a solid [52], see Appendix for details.

With increasing *p*_0_, the storage shear modulus decreases at all frequencies, while the loss shear modulus declines only at higher frequencies. Tong et al. [18] demonstrated that in this regime, the elastic constants *k*_1_ and *k*_2_ of the Zener model (see Appendix) linearly decrease with *p*_0_ and reach zero at the solid-fluid transition when 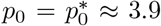. In contrast, the dashpot constant *η*_1_ remains mostly unaffected by changes in *p*_0_ and is primarily influenced by the friction parameter *ξ*, which is the sole dissipation factor. As a result, the characteristic time scale *η*_1_*/E*_1_ diverges following 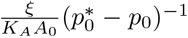 as *p*_0_ approaches the transition, determining the crossover frequency between these two regimes.

In the fluid-like phase where the zero-strain-rate modulus *G* is zero and *p*_0_ ≳ 3.9, *G*′ and *G*″ exhibit different behavior. The loss modulus, *G*″, predominantly scales linearly with frequency, *G*″∝ *ω*, and becomes dominant at lower frequencies. Conversely, the storage modulus, *G*′, exhibits a quadratic frequency dependence, *G*′∝ *ω*^2^, and approaches zero as frequency decreases to zero, indicative of fluid-like behavior. This matches the Standard Linear Liquid or Burgers model in rheology, which consists of two Maxwell element in parallel (Appendix).

### B. Confluent Active Foam Model

We seek to quantify the stability of the confluent active foam model as a function of the interfacial tension parameter, Λ, which exactly follows the analysis of Staple et al [47] for the full standard vertex model, except that here we drop the *p*^2^ term. First, we nondimensionalize Eq. 13 for a single cell to get 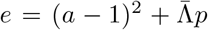 where 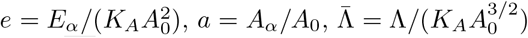 and 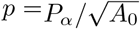 To look for energy minima, we scale each edge of the polygon by 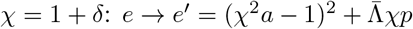 and require ∂*e*′/∂χ|_χ=1_ = 0. We find

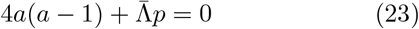

Two trivial solutions are i) *a* = 0, *p* = 0 corresponding to a collapsed polygon, and ii) *a* = 1, *p* = 0, which is not possible to achieve, suggesting that the polygons are frustrated at the energy minima. A nontrivial solution occurs when a < 1 (otherwise ∂*e*′/∂χ|_χ=1_ > 0).

Looking at 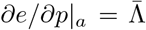, we see that for 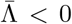, the energy can be reduced by increasing *p*, which makes the polygon unstable to shear. For 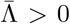 to reduce energy we need to reduce *p* as well, suggesting that a regular polygon is the solution.

For a regular polygon, we write *a* = *cp*^2^ with *c* = cot(π/*n*)/4*n* and *n* the number of edges. Then we can rewrite 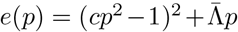 and look for the solution to 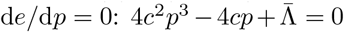. Additionally, we require the polygons to have *p* > 0 (not collapsed):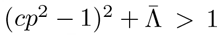. This suggests that there is a range of 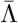 that supports a stable (rigid) polygon:

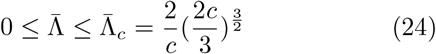

This range is exact only for regular polygons, so in simulations of disordered cellular tilings, we expect that the observed transition point 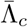 deviates from this ordered prediction. This is indeed what we find; in disordered vertex model simulations some cells collapse for Λ greater than a threshold value (shown in panel c of Fig 2), while for Λ < 0 the simulation is unstable since there is no restoring force and the energy can keep decreasing for larger and larger perimeters. In this case, vertices cross into neighboring cells and the model becomes unphysical, as illustrated in Fig 2(a).

**FIG. 2.**
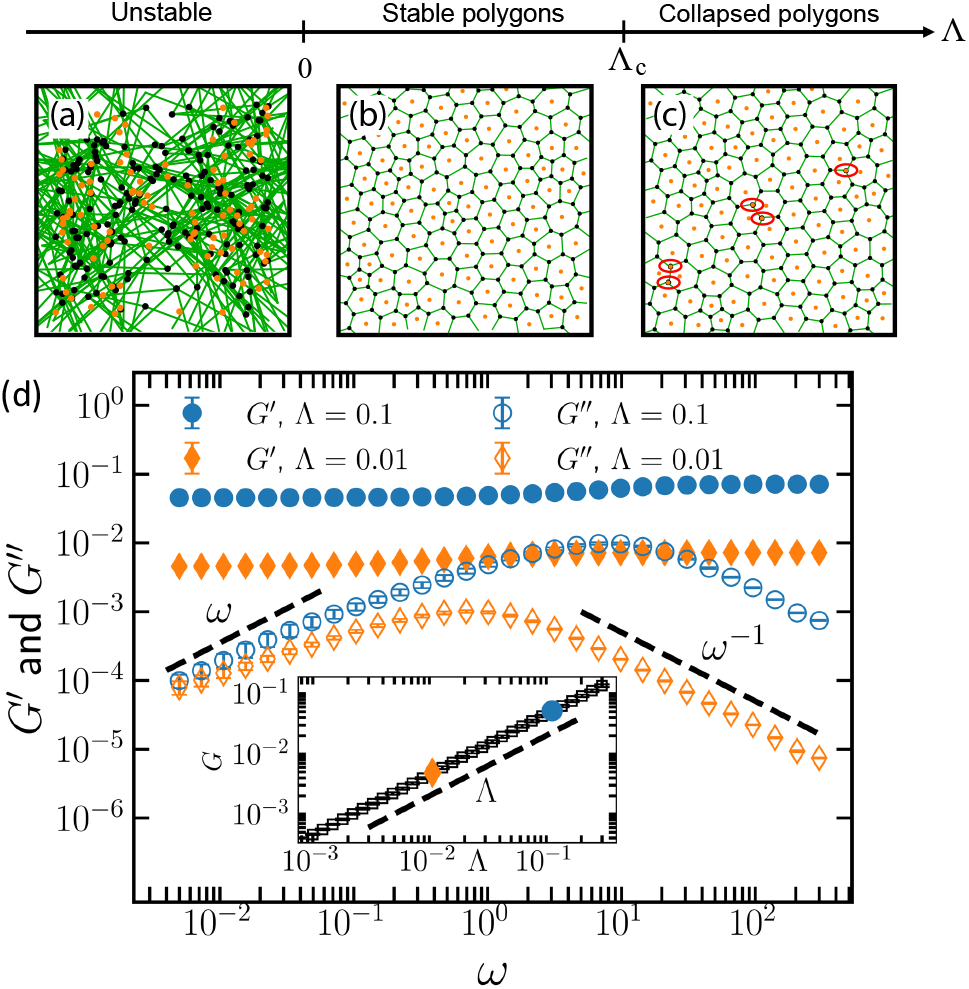
Stability and Mechanical response of the Confluent Active Foam Model. (a-c) Snapshots from numerical simulations of disordered tissues at different values of the interfacial tension Λ. (a) Snapshot during an unstable simulation for Λ < 0, illustrating a runaway instability where cell edge lengths increase indefinitely. (b) Stable steady state of simulation with 0 < Λ < Λ_*c*_. (c) Early stage in a simulation with Λ > Λc where cells continue to shrink, leading to an indefinite coarsening of the foam. In (c), a few cells have already collapsed in the middle and lower left-hand corner, indicated by their centers—marked with orange dots—resting on top of vertices and highlighted by a red ellipse. (d)Frequency dependence of the storage and loss moduli, *G*′ and *G*′′, within the stable regime 0 < Λ < Λ_*c*_ for two selected values of Λ, highlighted in the inset. Inset: log-log plot of the zerofrequency shear modulus *G* as a function of interface tension Λ for 0 < Λ < Λ_*c*_. The dashed line, with a slope of 1, illustrates the linear relationship between *G* and Λ.

This implies that the confluent limit of the active foam model is incapable of exhibiting the fluid-like phase observed in standard vertex models, as any potential rigidity transition would occur at Λ < 0, a range where the model becomes unstable. The inset to Fig. 2(d) is a loglog plot illustrating that the zero-frequency shear modulus *G* increases linearly with the tension parameter Λ, and vanishes in the limit Λ → 0.

To analyze the rheological properties of this model in the region it is stable, we performed frequency sweeps using dynamic oscillatory techniques. The results, illustrated in Fig. 2(d), reveal that the storage modulus, G′, exhibits a plateau, while the loss modulus, *G*′′, scales linearly with frequency at lower frequencies, indicating solid-like phase. At higher frequencies, *G*′ reaches a higher stable value, and *G*′′ scales as *ω*−1. These results align well with the predictions of the Zener model, and confirm that the active foam model has the same rheological scaling as the standard vertex model in its solid phase.

### C. Spring Edge Model

We next discuss the spring edge model, where every edge is governed by its own linear spring of rest length l0. First-order rigidity theory, also known as Maxwell constraint counting [31], suggests that the model should never be floppy: *N*_*d*.*o*.*f*_. = 4*N*_*cell*_, *N*_*constraints*_ = *M* = *N*_*cell*_ + 3*N*_*cell*_ = 4*N*_*cell*_, therefore *N*_*d*.*o*.*f*_. = *M*. A similar situation occurs in the 2D Voronoi model, as discussed by Sussman and Merkel [38], and also in fiber networks with bending moduli [53].

Even though such systems do not exhibit a singular rigidity transition, there is still a crossover from soft to stiff behavior. In the 2D Voronoi model, no rigidity transition is observed in the limit of zero temperature (activity), but in the presence of finite fluctuations there is a strong crossover from solid-like behavior to fluid-like behavior at a critical value of *p*_0_ [38]. Similarly, fiber network models with a bending energy are always rigid, but the network can be arbitrarily soft at small strains and connectivities, and the stiffness increases dramatically as the connectivity or strain increases past a critical value.

As highlighted in Figure 3, simulations at zero temperature confirm that Maxwell constraint counting is predictive for the spring edge model; all configurations have a finite shear modulus. However, we observe that the system exhibits a crossover from very soft to stiff as the model parameter *l*_0_ decreases. This crossover involves a change in cell shapes from more regular and convex to more irregular and concave. The crossover becomes sharper as *K*_*A*_ is decreased and in the limit of *K*_*A*_ = 0 it becomes a singular rigidity transition (Fig 3(a)), as has been observed previously for spring networks [44, 54, 55], and the drop in the magnitude of the modulus at the crossover for *K*_*A*_ ≠ 0 scales with *K*_*A*_, as highlighted in the inset to Fig 3(a). This is highly reminiscent of observations in spring networks with a bending energy, and suggests that the area term in the spring-edge model is playing the same role as the bending term in fiber networks.

**FIG. 3.**
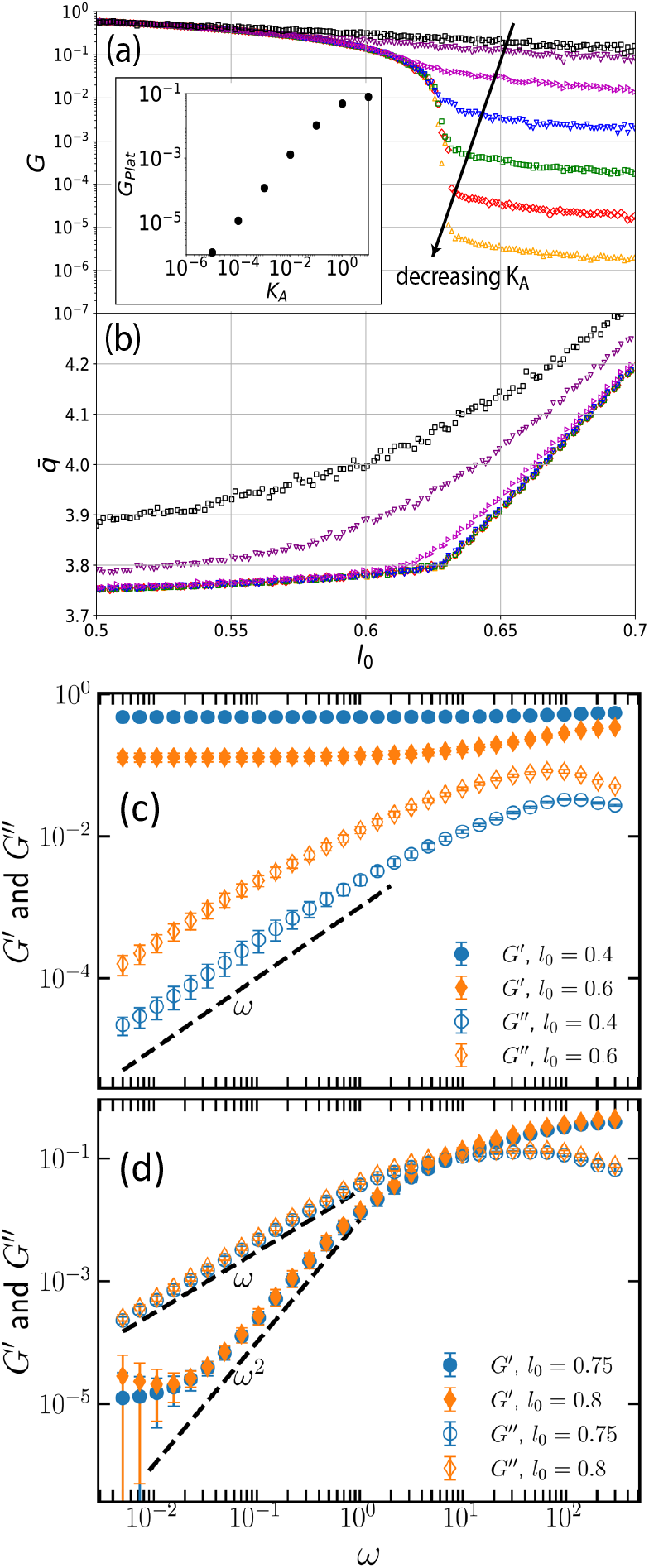
Mechanical response of the Spring-Edge Model. (a) Zero-strain-rate shear modulus *G* as a function of the spring rest length *l*_0_ across various values of area stiffness *K*_*A*_ ∈ (10^−5^, 10^−4^, 10^−3^, 10^−2^, 10^−1^, 1, 10) in the Spring-Edge model. Inset displays the plateau of the shear modulus *G*_*plat*_ within the floppy regime (large *l*_0_) as a function of *K*_*A*_. (b) Observed cell shape index as a function of l0 for the same range of *K*_*A*_, demonstrating a significant change in behavior at the crossover point, akin to that observed in the standard vertex model. (c) Frequency dependence of the storage (G′) and loss (G′′) moduli in the solid phase for two different values of the rest length, as indicated in the legend. (d) *G*′ and *G*′′ as a function of frequency in the fluid phase for two different values of the rest length, as shown in the legend. These viscoelastic results in (c) and (d) correspond to a *K*_*A*_ value of 10^−5^.

These observations suggest that when the area stiffness *K*_*A*_ is set to small or intermediate values, the spring-edge model demonstrates behavior similar to the standard vertex model. Specifically, there is a transition from a nearly floppy state characterized by a low shear modulus to a stiff state with a high shear modulus as the cell interfaces become stiffer (e.g., as *l*_0_ decreases).

Previous work on the standard vertex model highlights a strong connection between observed cell shape and the floppy-to-rigid transition, which is also observed in experiments [5, 6]. Therefore, an obvious question is whether there is a similar relationship in the spring-edge model. Fig 3(b) shows the observed 2D shape index 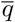 as a function of model parameter *l*_0_ for different values of *K*_*A*_. In the parameter regime where the tissue is stiff, 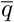 changes very little and is pinned near a characteristic value 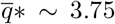, while in the parameter regime where the tissue is nearly floppy, 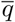 rises quickly away from that characteristic value. Qualitatively, this behavior is nearly identical to what is observed in standard vertex model simulations [5, 6], and predicted by vertex model scaling arguments [44].

Taken together, this highlights that at least in some parameter regimes, the spring-edge model exhibits shape-tissue fluidity correlations that are remarkably similar to those in standard vertex models. This in turn suggests both that i) an observation of such a correlation in experimental data can not be used to distinguish between models and ii) a strong correlation between shape and tissue fluidity could potentially be a universal feature of a broad class of models.

To study whether the finite-strain behavior also exhibits universality, we analyzed the finite frequency behavior. Figure 3(c) and (d) shows the storage and loss moduli of the spring-edge model at *K*_*A*_ = 10−5. For *l*_0_ ≲ 0.65 the system behaves as a solid and we observe that the storage modulus exhibits a plateau while the loss modulus scales as *G*′′ ∝ *ω* at low frequencies. At high frequencies, *G*′ attains a higher plateau but *G*′′ decreases with frequency. These scalings and trends are identical to the viscoelastic behavior of the standard vertex model in the solid-like shape index regime. As the rest length *l*_0_ increases, the storage modulus decreases across all frequencies, mirroring behavior observed in the standard vertex model. However, the loss shear modulus *G*′′ exhibits a distinct response, increasing at all frequencies, which is different from the patterns seen in the standard vertex model as *p*_0_ increases.

On the other hand, for *l*_0_ ≳ 0.65, *G*′′ exceeds *G*′ over a wide range of low frequencies, indicating a fluid-like response (Fig. 3(d)) in this regime. For that range of frequencies, 10−2 ≲ *ω* ≲ 1, G′ ∝ *ω*^2^ and *G*′′ ∝ *ω*, consistent with scaling behavior seen in the standard vertex model in the fluid-like phase. At the very lowest frequencies, the storage modulus *G*′ exhibits a plateau, indicating that *G*′ will eventually dominate over *G*′′, consistent with the very small but finite zero-strain-rate shear modulus shown in (a). We observe that this low-frequency crossover shifts to higher values as the area stiffness *K*_*A*_ increases. In summary, the finite-strain-rate rheology of the spring-edge model also shares remarkable similarities with the standard vertex model, including the same rheological scaling behavior over a wide range of frequencies.

### D. Dynamical models

We will first study whether dynamical models also exhibit a rigidity transition in the limit of infinitely slow driving. To calculate the standard definition of the shear modulus in presence of zero fluctuations, a model must exhibit a force-balanced steady state in that quasi-static limit; otherwise, the steady state will be dynamic with timescales that compete with the shear driving rate, and the shear modulus is not well-defined.

In what follows, we distinguish between the vertex degrees of freedom {*r*_*i*_} (or equivalently, {*A*_*α*_, {*l*_*ij*_}) and the internal degrees of freedom, such as the rest lengths 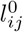 and myosin density *m*_*ij*_ for the active tension model.

If a model’s system of differential equations does possess a steady state where all the vertices {*r*_*i*_} remain at fixed positions that correspond to a local energy minimum, we term that a quasi-static steady state (QSS). Obviously many dynamical systems do not have such a state. Therefore, to see if such a state exists, we first construct nullclines of the dynamical model to identify possible steady states. Next, we attempt to construct a static vertex energy function (denoted by *E*^*^) restricted to the steady state configurations, where, for example, the tensions in Eq 1 are no longer functions of time but only vertex coordinates. Finally, we analyze whether that static system defined by *E*^*^ exhibits a response to vertex perturbations that is equivalent to dynamics of the full dynamical system near that fixed point. If so, we have successfully identified a QSS with a well-defined shear modulus.

First, we argue that in the limit of fast vertex dynamics and slow internal DOF dynamics none of these dynamical models possess a QSS on disordered networks. To see this, assume that vertex relaxation happens at a fast rate, so that before internal states have changed much, vertices have reached mechanical equilibrium. This equilibrium generically involves a distribution of tensions for random packings. Thus at timescales when the internal dynamics start to become relevant, each rest length will change by a different rate based on the tension on the edge. This will break force balance and cause vertices to move again, negating the existence of a steady state. In other words, the fast vertex dynamics are always coupled to the slow internal rest length dynamics and a QSS is not achieved.

Of course the system can also achieve a steady state when the dynamics of both sets of degrees of freedom simultaneously reach their fixed points, but such a system will exhibit dynamics over finite timescales and also does not have a QSS. Therefore, in what follows we focus on the limit that the internal dynamics are fast compared to vertex degrees of freedom.

#### 1. Active Spring Model

For the active spring model, first assume that *ϵ*_*c*_ = 0. Then at steady state, 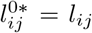 and 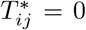. Thus, the equivalent static energy *E*^*^ is given by

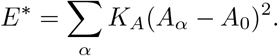

More generally, for *ϵ*_*c*_ > 0, some edges will still hold tensions but the maximum strain on an edge will be *ϵ*_*c*_. In other words, for those edges with |*ϵ*_*ij*_(t = 0)| > *ϵ*_*c*_ the steady state will be reached when 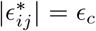 or 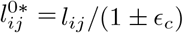:

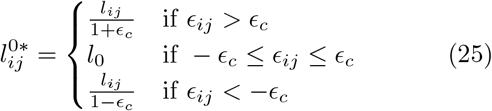

where *l*_0_ is the initial rest length of the whole system. The tension *T*^*^ at steady state is

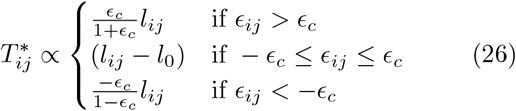

To get the equivalent static energy *E*^*^, we insert the tensions in Eq. 1 and integrate. The reason we should not use the spring edge energy function is because in this steady state system, rest length is a function of *l*_*ij*_ and so needs to be properly integrated.

One can use this equivalent static system to show (see Appendix B1) that for the Active Spring Model, the zero-strain rate limit only exists when *ϵ*_*c*_ = 0. In other words, only when *ϵ*_*c*_ = 0 is the linear response to perturbations of coordinates for *E*^*^ is equivalent to the dynamical model in the QSS limit.

Next, we seek to understand the finite-frequency response of the active spring model. We carried out simulations on random vertex networks with equations of motion given by Eq. 10 and Eq. 16. Our results con firm that a stable fixed point for the energy function is generally unattainable except when *ϵ*_*c*_ = 0. In the case of *ϵ*_*c*_ = 0, the system exhibits nonlinear, transient dynamics (See Appendix Fig D2). Therefore, we explored the viscoelastic properties of this model by analyzing the storage and loss moduli, *G*′ and *G*′′, across various frequencies *ω* and waiting times *t*_*w*_ (Fig. 4(a-b)) as described in Section IIA3.

**FIG. 4.**
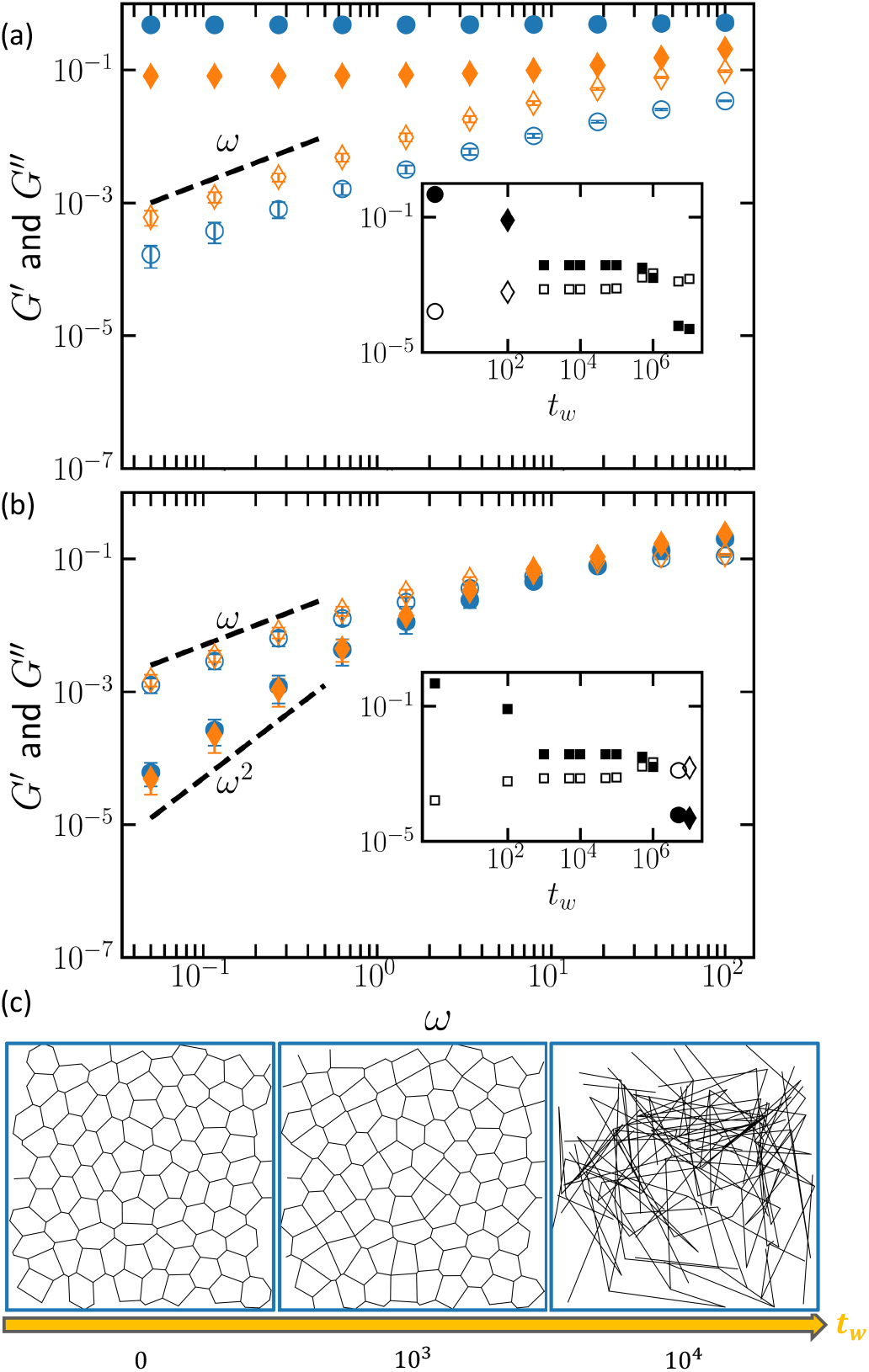
(a,b) Mechanical response of the Active Spring-Edge Model. (a) The behavior of the storage and loss moduli, *G*′ and *G*′′, for the Active Spring Model as a function of oscillation frequency *ω* in the solid-like phase (small waiting time). The two distinct set of data are for two different *t*_*w*_ as highlighted in the inset. The inset presents *G*′ (closed symbols) and *G*′′ (open symbols) plotted against *t*_*w*_ at a low frequency (*ω* = 0.05), illustrating the fluid-like behavior of this model at intermediate *t*_*w*_. (b) The behavior of *G*′ and *G*′′ as a function of oscillation frequency *ω* in the fluid-like phase (long waiting time). The area stiffness is 10−5 here. (c) **Stability of Tension Remodeling Model**. Snapshots from simulations of the Tension Remodeling Model at different timepoints *t*_*w*_ from the onset of tension remodeling dynamics at t=0, illustrating that an instability generically occurs at long enough timescales.

Figure 4 illustrates the behaviors of *G*′ and *G*′′ for this model initialized with a uniform rest length *l*_0_ = 0.4. At shorter waiting times *t*_*w*_, the system behaves similarly to the conventional spring-edge model and standard vertex model, displaying a solid-like response. Here, *G*′ reaches a plateau and *G*′′ linearly approaches zero as the frequency decreases. Conversely, as *t*_*w*_ is increased, the edge tensions reduce, rendering the system more fluidlike. This transition is clearly evidenced in Figure 4 through the corresponding changes in *G*′ and *G*′′, where *G*′ decreases and *G*′′ becomes dominant at lower frequencies.

In summary, the active spring and tension remodeling models does not possess a quasi-static steady state except in the limit of *ϵ*_*c*_ = 0, where the network is floppy. Therefore, it is not possible to calculate a shear modulus independent of the applied shear rate for this models, and it is clearly distinct from the standard vertex model in the limit of slow shear. At finite strain rates, after short waiting times displays a solid-like rheology similar to the standard vertex model, but in the long waiting time limit internal dynamics drive the systems towards a fluid-like state that is distinct from standard vertex model, consistent with the zero-strain rate analytic results.

#### 2. Tension Remodeling Model

For the tension remodeling model, assume that we start with uniform tension *T*_*ij*_(t = 0) = Λ on all edges. At steady state,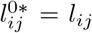. If |*ϵ*_*ij*_(t = 0)| > *ϵ*_*c*_, we have 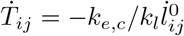. Integrating this gives

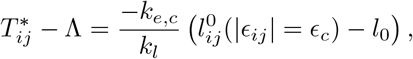

which leads to

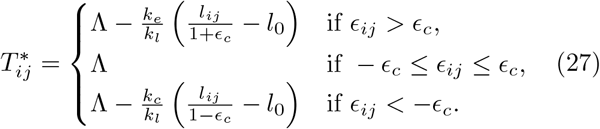

The tension can now show spring-like behavior. As before, integrating Eq. 1 with 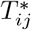 gives us the equivalent static model energy, *E*^*^.

One can show (see Appendix B2) that tension remodeling only has an equivalent QSS in the limit of *ϵ*_*c*_ = 0. Even then, the equivalent static model does not have a stable fixed point in simulations, which can be predicted from Eq. 27: the spring strains on the edges have the wrong sign and do not act as restoring forces. Simulation snapshots highlighting the unstable nature of this model are shown in Fig. 4(c). Due to these serious pathologies, it is not possible to compute a shear modulus, and we do not study or report the finite-frequency response of the tension remodeling model.

#### 3. Active Tension Network

For the active tension model, at steady state the tension is dictated by the myosin distribution: 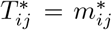 Moreover, from Eq. 22, 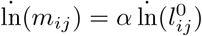 so that

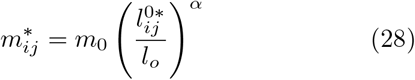

where we have assumed that a uniform myosin field *m*_0_ at *t* = 0. If we further assume 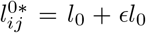 such that |ϵ| ≪ 1 we can simplify Eq. 28 to read

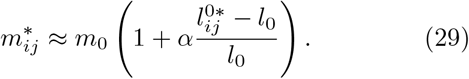

From 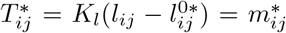 we can then describe 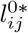 in terms of *l*_*ij*_.

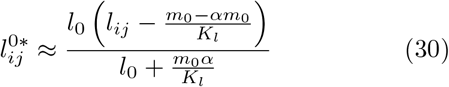

Finally, the tension is given by

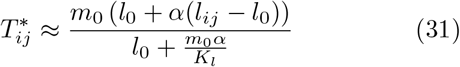

which, similar to the previous models, exhibits spring-like behavior. In fact, we can rewrite Eq. 31 as

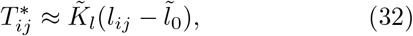

where the effective rest length is

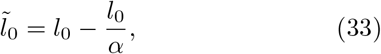

and the effective spring constant is

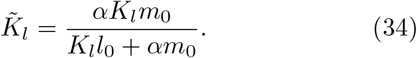

This equation indicates that the two myosin activity parameters *m*_0_ and *α* allow the effective rest length and effective spring constant to be independently varied.

One can show (Appendix B3) that unlike the other two dynamical models, the active tension network matches the equivalent static model in the limit of fast internal dynamics. We have confirmed in numerical simulations that a system with tensions given by Eq. 31 is indeed able to find a stable energy minimum.

Next, we calculate the shear modulus in simulations of this vertex model. Here we present data for *K*_*A*_ = 0.0, but the data is quite similar for small *K*_*A*_ > 0.0. Fig 5(a) illustrates the ensemble-averaged shear modulus *G* as a function of the model parameters *l*_0_ and *α*. The contours illustrate lines of constant effective rest length 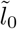 (Eq. 33). For example, at 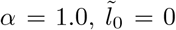 for any *l*_0_. Above this line corresponds to 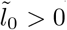 while below is 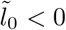 For large values of the spring rest length and myosin recruitment rate, the shear modulus is zero and the system is floppy, while for lower values of these parameters the system becomes rigid.

**FIG. 5.**
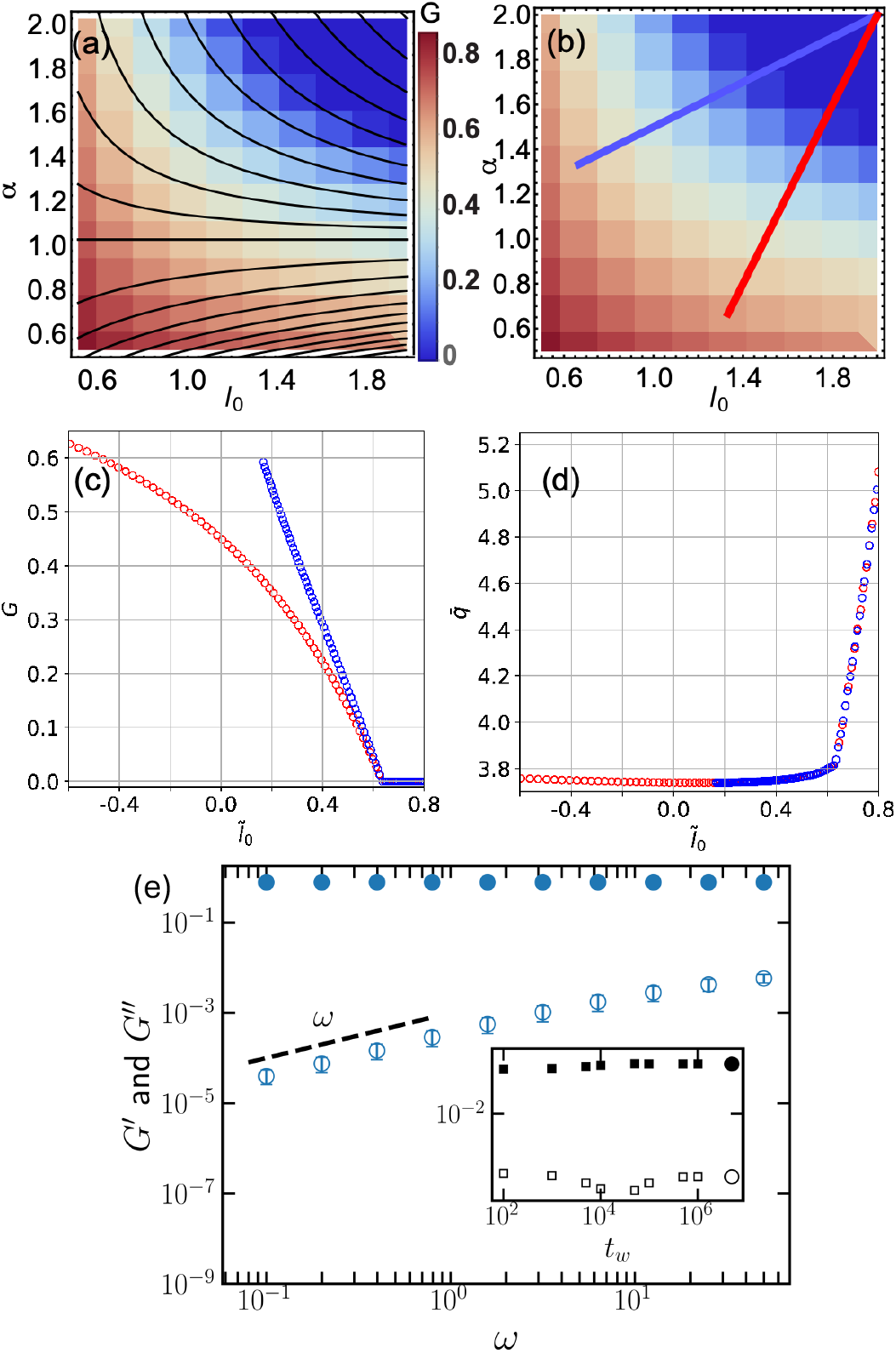
Mechanical Response of the Active Tension Model. (a) Shear modulus of the quasi-static active tension model as a function of myosin recruitment rate α and spring rest length *l*_0_. Contours illustrate lines of constant effective rest length 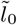 (Eq. 33). (b) Same plot as figure (a), where red and blue lines illustrate two trajectories with linearly increasing effective rest length 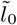. (c) Shear modulus as a function of effective spring rest length 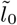 for two different trajectories through model space, illustrated by the red and blue lines in panel (b). (d) Observed shape index 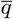 as a function of effective spring rest length 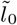 for two different trajectories, illustrating a strong crossover in shape at the location of the floppy-to-rigid transition. (e) The storage and the loss moduli, *G*′ and *G*′′ as a function of frequency at a long waiting time highlighted in the inset. For this data, the tensions are initialized at 0.05 and *α* = 10−5. The inset shows *G*′ (closed symbols) and *G*′′ (open symbols) as a function of waiting time *t*_*w*_ at a low frequency *ω* = 0.1.

**FIG. 6.**
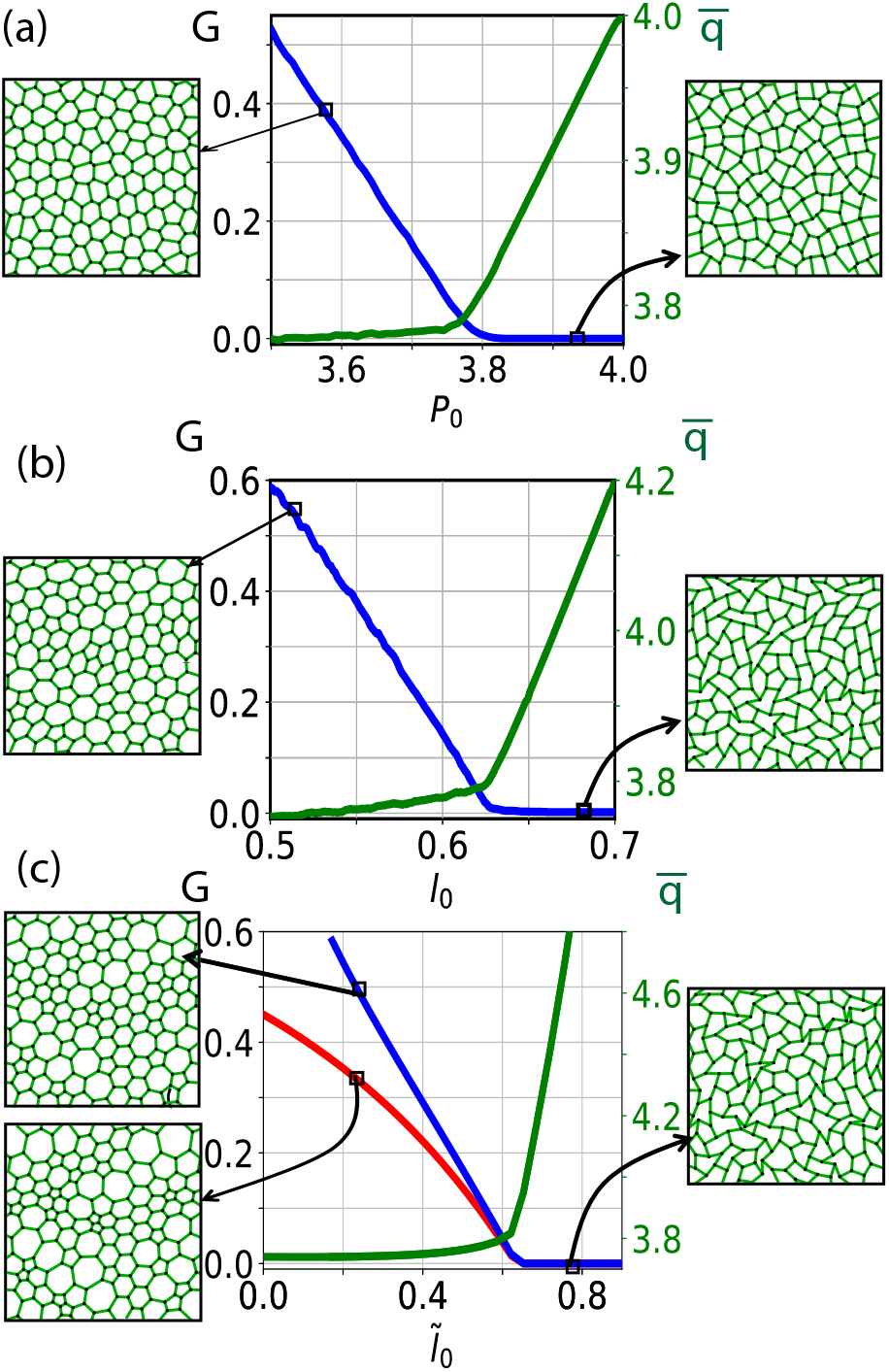
Schematic diagram of universal behavior in vertex models. All three models, the (a) Standard vertex model, the (b) Spring-Edge model, and the (c) Active tension model exhibit a crossover from soft (low shear modulus *G*) to stiff (high shear modulus) behavior as a function of the control parameter, with an associated change in observed cell shape 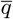.

To understand the origin of this behavior, we study the linear response along lines of linearly increasing effective rest length 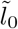 (Eq. 33), illustrated by the blue and red lines in Fig 5(b). Fig 5(c) demonstrates that as a function of the effective rest length, the active tension network is quite similar to the spring-edge model, with a shear modulus that is zero above a critical value of the effective rest length 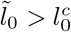.

Importantly, the extra model parameter *α* generates extra flexibility in the model, modulating the effective spring constant. Therefore, although the rigidity transition always occurs at the same value of the effective spring constant, the stiffness of the system once it becomes rigid can be independently regulated. Additionally, the model can exhibit 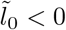, which normally would be unphysical for a spring network but here it can be achieved with *l*_0_ > 0 and *α* < 1. This also allows the system to become stiffer than in a simple spring-edge model.

Fig 5(d) shows the observed cell shape index 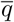 as a function of 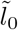 for the same two trajectories in parameter space highlighted in panel Fig 5(b). Importantly, the shape parameter shows the same behavior as all other variations of the vertex model with a well-defined shear modulus that we have studied: 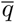 remains pinned at a low value of 3.75 in the stiff/rigid phase of the model, and rises sharply away from that value in the floppy regime.

We examined the finite frequency response of this model, with results shown in Figure 5(e). We initialized the systems with parameters for rest lengths, tensions, myosin, and *α* that locate the model in the solid-like regime. After that, we turned on the internal dynamics for all these degrees of freedom. Allowing these dynamics to continue, we observe the breathing mode known as *isogonal* transformation [3]. We then find the minimum energy state of the system after a waiting time *t*_*w*_ and perform the standard oscillatory shear tests to calculate *G*′ and *G*″. As shown in Fig. 5(e), we find a solid-like response in which *G*″ shows a linear scaling with frequency. This is similar to the behavior of a static spring-edge model at low *l*_0_. To transition into a fluidlike state, increasing *l*_0_ is necessary, as indicated in the zero-frequency phase diagram of Fig. 5(a). However, the computational model becomes unstable for *l*_0_ > 1.0. Using a different method, Noll et al. [3] showed that this model becomes fluid-like in the low frequency limit without freezing the internal degrees of freedom by using an extensional rheology method and computing the phase shift.

## IV. DISCUSSION AND CONCLUSIONS

We have studied the rigidity transition and viscoelastic response in six different variations on the vertex model for confluent tissues. We find that a subset of them – the spring-edge model and the active tension network – exhibit a crossover from floppy to stiff at a characteristic value of a model parameter: the effective spring rest length, accompanied by an abrupt change in the behavior of the observed cell shape, when *K*_*A*_ is small. Moreover, these models all exhibit the similar scaling of storage and loss modulus with frequency, consistent with the behavior of the Zener solid on the rigid side of the transition and a Burgers fluid on the fluid side of the transition.

What about the other models? The confluent foam model is numerically unstable in the regime that would be floppy, and so there is no floppy-to-rigid or fluid-to-solid transition in that model, though the rheology is the same as the standard vertex model on the solid side of the transition.

The other two *dynamical models* we studied do not have a well-defined quasi-static limit. All versions of the tension remodeling model, as well as the active spring model with *ϵ*_*c*_ ≠ 0, are pathological in the large-time limit due to the generic emergence of an instability. This suggests they may not be useful for describing biological tissues in the limit of long timescales.

The active spring model with *ϵ*_*c*_ = 0 is well-behaved, but always dynamic, even in the limit of very long times/slow strain rates. This means it is not possible to formally define a shear modulus, and the viscoelastic response is highly nonlinear due to changes to spring rest lengths over time, depending on the waiting time since the system was initialized. We find that the model is solid-like during initial transients and becomes fluid-like on long timescales, with the same frequency scaling as the standard vertex model.

To summarize, in parameter regimes where vertex models are not pathological, they have floppy-to-rigid crossovers predicted by the same simple observable, the cell shape parameter. They also have the same scaling of their finite-frequency viscoelastic response on either side of the transition. This suggests that there is a broad class of energy functionals (some even with extra dynamics) that all generate the same universal features, and helps to explain why the cell shape parameter has been so successful and useful in experiments.

An important point is that here we have only demonstrated universal behavior in some aspects of the response, and it would be useful and interesting to search for additional features in the future. For example, the gold standard for claiming universality in physics problems is to demonstrate that the critical scaling exponents for the transition are the same across multiple systems. Unfortunately, due to the cusp-y nature of the energy landscape in vertex models [39, 56] and large finite-size effects near the critical points in spring networks [44, 57], numerical minimization and computation of shear modulus near transition points pose significant challenges [10, 44]. This makes interpreting critical scaling exponents difficult and subtle, leading to ongoing discussions within the community [44, 55, 58]. Moreover, for several of the models we study here, the change in rigidity will be a crossover instead of a *bona fide* transition. Nevertheless, it would be exciting to perform a crossover scaling analysis, e.g. similar to those performed other systems with a crossover from one type of rigidity to another as in Ref. [59]), and search for universal exponents.

Not everything is universal; there are also some interesting differences between models. For example, the extra myosin degrees of freedom introduced in the active tension network (which were needed to match experimental observations) give that model a higher degree of tunability than the standard vertex model. Simply by tuning the myosin recruitment parameter *α*, the tissue is able to tune its collective overall stiffness, which could be functionally beneficial in developing embryos. It would be interesting to investigate how various fruit fly mutations alter this parameter and study whether the observations of global embryo shape and dynamics are consistent with changing overall stiffness.

This raises the question of why the cell shape is a good order parameter for the rigidity transition in so many models, since the control parameter in both the spring-edge and active tension models is not the shape, but the spring rest length. One hint is provided by recent work that indicates a geometric incompatibility [44, 45] is driving a second-order rigidity transition in the standard vertex model, and that the same mechanism drives strain-stiffening in fiber networks [31]. These works highlight that there are two length scales in the system: 1) one defined by the number of cells/vertices per unit area and 2) the characteristic distance between two cells or vertices defined by the energy functional (parameterized by the cell perimeter in vertex models or the rest length for edges in spring-edge models).

Rigidity occurs when the second, energy-defined length scale becomes smaller than the first, density-based length scale. In the rigid phase, there appear to be no states with the specified density that allow the system to have zero energy, and the second-order rigidity transition occurs at a special point in configuration space where states that are compatible with both the energy length scale and the density length scale disappear. Given that all these models seem to have a second-order rigidity transition, it could be that the observed cell shape index, which is one of the obvious dimensionless numbers that compares those two length scales, is a good order parameter for the transition across all types of models for tissues. More work is needed to understand whether many (or all) second-order rigidity transitions share these universal geometric features, which could allow us to make these ideas quantitatively precise.

A related question is why vertex models seem to be such an effective representation of the degrees of freedom for understanding rigidity transitions in epithelial tissues. For example, we could instead have chosen to represent the degrees of freedom as in a Cellular Potts Model [60], where each cell is composed of tens to hundreds of grid points that represent the cell body. Such models also seem to have a rigidity transition controlled by cell shape [61]. Or we could have gone to even smaller scales to represent features of the actin cytoskeleton or adhesion molecules.

One possible answer is suggested by the previous paragraphs: if the fundamental cause of the rigidity transition is an incompatibility between two length scales (an internal length scale generated by the cytoskeleton that creates an energy penalty for changes away from a homeostatic set length, and a second length scale that characterizes the number density of cells in a container of a given size), then we may want to use the simplest model that correctly incorporates those two length scales and the tension between them. This turns the typical physics approach on its head; normally we first define our degrees of freedom and their interactions and then study the emergent behavior. In some sense, this analysis suggests that once we understand the universal physics of the rigidity transition, we can use it to identify the right scale for coarse-graining (i.e. how to best define the degrees of freedom). One wonders if other problems in biology might lend themselves to a similar approach.

## V. ACKNOWLEDGEMENTS

We thank Emanuela Del Gado and Suzanne Fielding for insightful discussions. We acknowledge support from the Simons Foundation grants No 454947 to MLM (OKD and MLM) and No 446222 (MLM and ELK). SA and MLM acknowledge support from the Chan Zuckerberg Theory in Biology initiative, and MLM acknowledges support from NSF-DMR-1951921 and NSF-CMMI-1334611.

## Appendix A: Stress tensor for vertex models

The stress tensor for a cell *c* is computed as in Refs. [62–64]

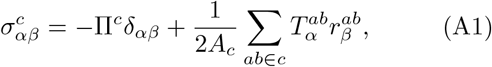

where *α* and *β* denote the Cartesian coordinates and *δ*_*αβ*_ is the Kronecker delta. The first term on the right-hand side is due to hydrostatic pressure inside cell with 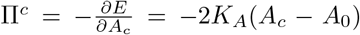. The second term on the right-hand side takes into account the stresses due to the tensions on edges *ab* of the cell *c*. Since each edge is shared by two cells, we need to use the factor 1*/*2. 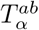 denotes the *α* component of the edge tension which is derived as 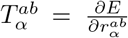, where 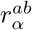 denotes the edge vectors between vertices *a* and *b* of an edge on the cell *c*. The tissue stress subsequently determined by averaging the cellular stresses weighted by their area, given as 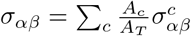, where *A*_*T*_ =Σ_*c*_ *A*_*c*_ is the total tissue area [64].

In this study, we explore various vertex models, each characterized by distinct energy formula for a cell *c*. The first category is the standard vertex model

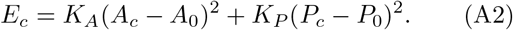

The second category includes models like the active foam model, which incorporate direct edge tensions

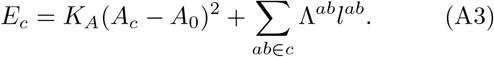

The third category uses springs on each edge, defined by a specific rest length

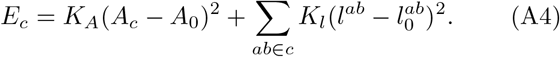

Although the hydrostatic pressure term Π^*c*^ is consistent across all models, the expression for the edge tension 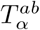, derived as 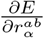, varies. For the first model, it is given by

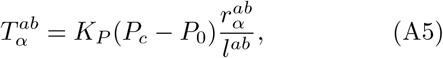

where *l*^*ab*^ denotes the length of edge *ab*. In the second model, the tension is

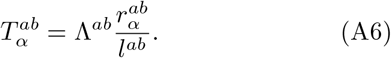

For the third model, the tension is

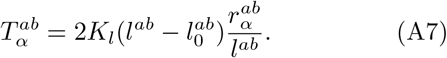

To study rheology of different vertex models under shear, we studied the shear stress component *σ*_*xy*_ under oscillatory simulations.

## Appendix B: Comparison of equivalent static models and full dynamical system for dynamic models

### 1. Active Spring Model

In the main text, we derived an equivalent static model for the Active Spring Model, Eq 26. Now, assume this equivalent static model has a stable fixed point. Does it have an equivalent dynamics to the actual dynamical system near the fixed point? In other words, if we perturb away from the fixed point 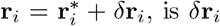 obtained from *E*^*^ the same as *δ***r**_*i*_ we get from the dynamical system? The answer should be yes if we are to claim *E*^*^ is an equivalent static representation of the dynamical model.

Here for simplicity, we assume the pressure is uniform across the tissue (or equivalently, *K*_*A*_ ≈ 0). Also, we assume that the edge strains |*ϵ*_*ij*_| > *ϵ*_*c*_ so that internal rest length dynamics has been activated (otherwise, the system would simply be a spring-edge model). Then, at equilibrium, 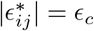.

First, some useful algebra: if 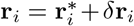, then to first order, 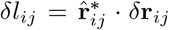, the unit vector along edge *l*_*ij*_ is 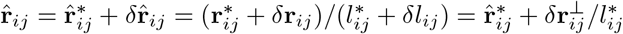 where 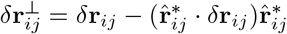.

The vertex dynamics is given 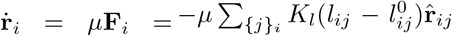.At equilibrium,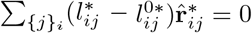. Then, for *δ***r**_*i*_ and 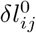 we have

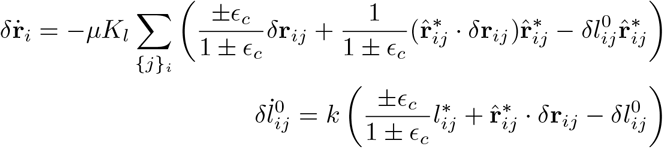

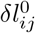 is only nonzero for edges with strains |*ϵ*_*ij*_| > *ϵ*_*c*_ after the perturbation. If we assume that, with a fast rest length dynamics compared to vertex relaxation, we get 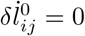 when |*ϵ*_*ij*_| = *ϵ*_*c*_, or

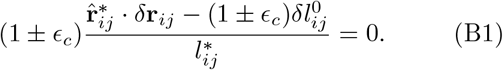

The solution is 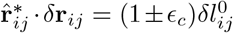. Plugging this back into the vertex relaxation dynamics, we get

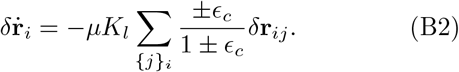

Now, let’s start from the equivalent static model. The tension is 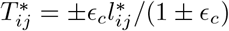. So

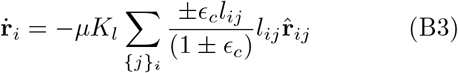

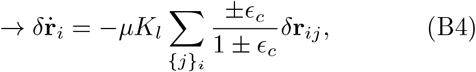

which is the same as the actual dynamical system. However, recall that Eq. B2 is only correct if |*ϵ*_*ij*_| > *ϵ*_*c*_, but that is not guaranteed generically. Almost surely, vertex perturbations will lead to some strains |*ϵ*_*ij*_| < *ϵ*_*c*_ in which case internal dynamics for those edges will not be activated and *E*^*^ will not generate dynamics equivalent to dynamical model.

This demonstrates that the zero-strain rate limit only exists when *ϵ*_*c*_ = 0. In other words, only when *ϵ*_*c*_ = 0 is the linear response to perturbations of coordinates for *E*^*^ is equivalent to the dynamical model in the QSS limit.

### 2. Tension Remodeling Model

Starting from Eq. 27 in the main text, we will follow the same procedure as the active spring model to see if the equivalent static model is equivalent to the QSS limit of the dynamical model in the limit of fast internal dynamics. Using force balance at equilibrium, 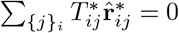, we find

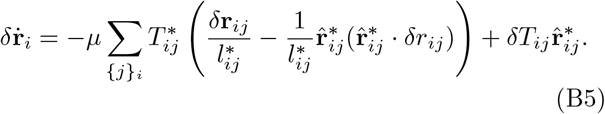

Here we assume that initially |*ϵ*_*ij*_| > *ϵ*_*c*_ so that the internal dynamics has been activated and 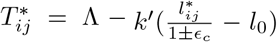 where *k*′ = *k*_*e,c*_ */k*_*l*_. Furthermore, we assume that the perturbation 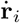 has led to |*ϵ*_*ij*_| > *ϵ*_*c*_ again so that *δT*_*ij*_ ≠ 0. From Eqs. 17 and 18 we can see

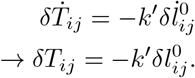

Note that *δT*_*ij*_(t = 0) = 0 and 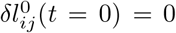. From Eq. 17, 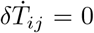 and QSS is achieved when |*ϵ*_*ij*_| = *ϵ*_*c*_ or 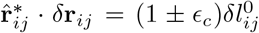 (Eq. B1). Thus at QSS we get 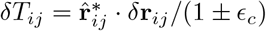 and

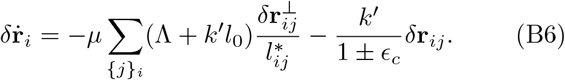

If we start with the equivalent static system, it is easy to see that we get the same vertex relaxation dynamics. However, again this is only true if *ϵ*_*ij*_ > *ϵ*_*c*_ for all edges after a perturbation, which is not generically the case. So, similarly to active spring model, tension remodeling only has an equivalent QSS in the limit of *ϵ*_*c*_ = 0.

### 3. Active Tension Network

For the Active Tension Network Model, we compare the behavior of the equivalent static energy functional perturbed from its minimum energy to the actual dynamical system near its fixed point. First, we focus on the equivalent static system. Starting from the tension given by Eq. 31, we can find *δT*_ij_ for the ES model in response to perturbation *δ***r**_*i*_:

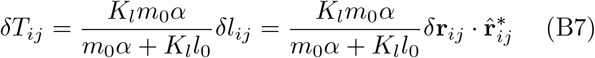

Now, we study the actual dynamical model. Vertex relaxation after perturbation is given by Eq. B5. We need to find *δT*_*ij*_ at QSS for comparison. To do so, we first write the 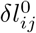 and *δm*_*ij*_ dynamics:

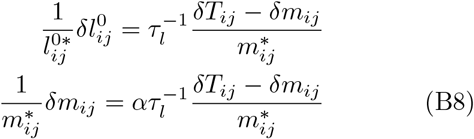

In the limit of fast internal dynamics then, we have *δT*_*ij*_ = *δm*_*ij*_. From Eq. 22 we also have

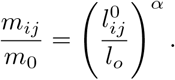

Assuming 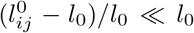 and using the fact that at equilibrium 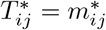 and *δT*_*ij*_ = *δm*_*ij*_, we find

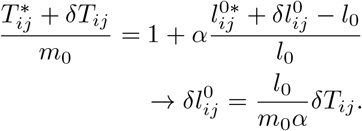

Since *T*_*ij*_ is the tension of a spring, we have

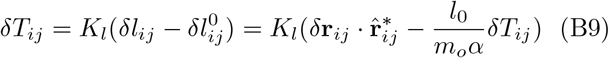

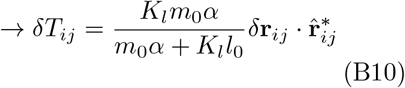

which is the same as *δT*_*ij*_ given from the effective static model, Eq. B7. Therefore, we can conclude that the active tension network matches the equivalent static model in the limit of fast internal dynamics.

## Appendix C: Oscillatory Rheology

To evaluate the storage modulus *G*′ and loss modulus *G*″ of viscoelastic materials within the linear regime, oscillatory rheology tests are performed. These tests involve applying a small amplitude oscillatory shear strain and measuring the resulting shear stress to calculate *G*′ and *G*″, as detailed in the main text. Figure C1 illustrates a schematic of such an oscillatory test. For dynamic models where internal degrees of freedom (DOF) evolve over time, we define a *waiting time* parameter *t*_*w*_. This parameter specifies the duration for which the dynamic model runs before the internal DOF are frozen to identify the minimum energy state for performing standard oscillatory tests.

**FIG. C1.**
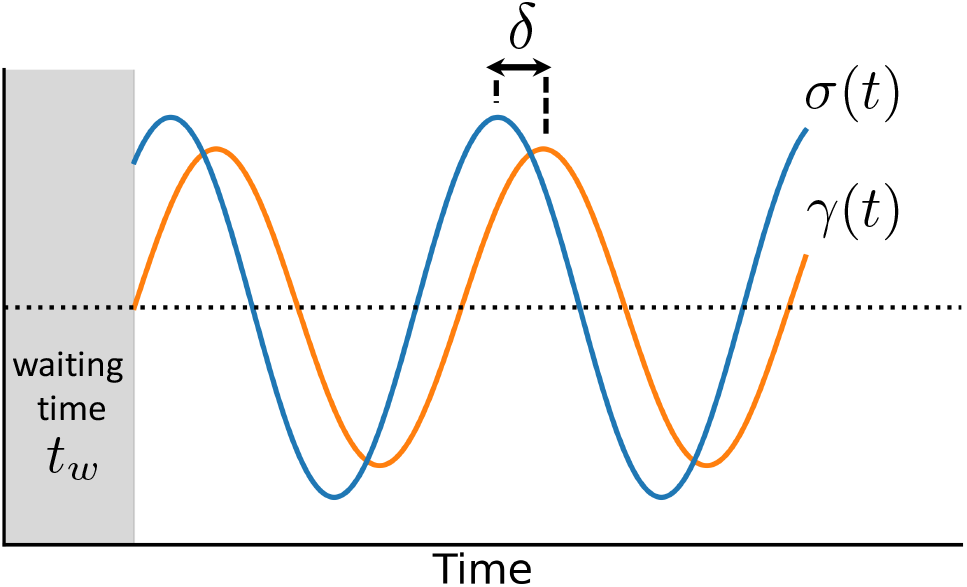
Schematic of Oscillatory Rheology Method. In oscillatory rheology, the system undergoes periodic deformation driven by a sinusoidal strain function at frequency *ω*. Within the linear response regime, the stress response is also oscillatory, exhibiting a phase shift *δ*. The complex modulus is calculated from the ratio of the Fourier-transformed stress to the strain signal, yielding the storage (real part) and loss (imaginary part) moduli. The shaded region shows the waiting time period. This is a parameter we use to explore rheology of dynamical vertex models. For static vertex models, *t*_*w*_ = 0.

## Appendix D: Lissajous–Bowditch Curves

A useful way to study the rheology of viscoelastic materials in an oscillatory test is to plot the stress *σ* as a function of strain *γ*. Eliminating time *t* from these properties within the linear viscoelastic regime yields an elliptical equation representing the stress-strain relationship.

These parametric plots, known as Lissajous–Bowditch curves, typically manifest as ellipses characterized by two symmetry axes—the major and minor axes of the ellipse. For a purely elastic material, following Hooke’s Law, this curve becomes a straight line at a 45° angle. In contrast, a purely viscous material shows a circular Lissajous–Bowditch curve. A viscoelastic material, exhibiting properties between these extremes, typically shows an elliptical Lissajous curve tilted at a 45° angle. Deviations from this elliptical shape suggest that the response has entered a nonlinear regime [65–67].

Figure D1 shows Lissajous–Bowditch curves for a standard vertex model. In the solid regime, the Lissajous–Bowditch curve becomes a line, in agreement with an system dominated by elasticity. Conversely, in the fluid regime, the curve forms a circle, characteristic of a viscosity-dominated system. As the frequency increases, the curve transitions into an ellipse, typical of viscoelastic behavior. The greater the elongation of the ellipse, the stronger the elastic response. As shown in the Fig. D1, after the first few oscillatory cycles, the system reaches a steady state behavior. When computing the storage and loss moduli, we remove the first 5 cycles.

**FIG. D1.**
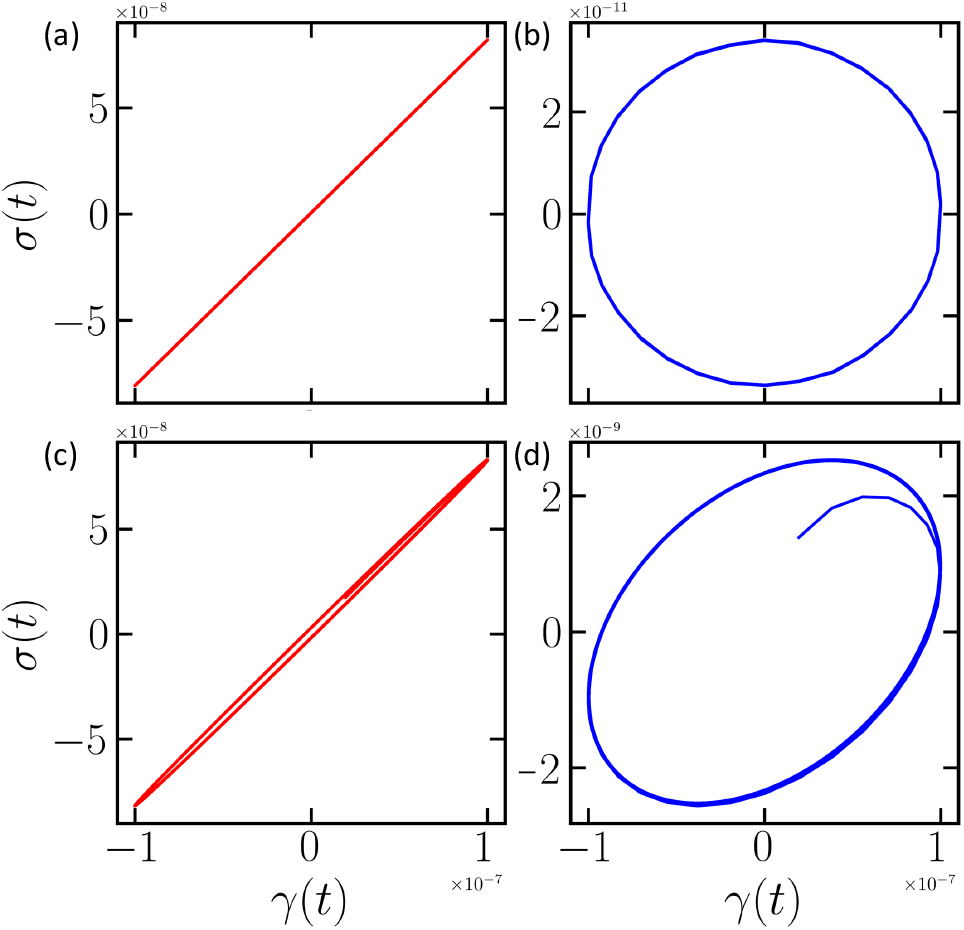
Lissajous-Bowditch curves for a Standard Vertex Model. (a) Solid-like regime with a target shape factor *p*_0_ = 3.4. (b) Fluid-like regime with a target shape factor *p*_0_ = 4.2. The oscillation frequency is *ω* = 0.01 here for a duration of 40 cycles. (c) and (d) panels shown the Lissajous-Bowditch curves at a higher frequency *ω* = 1.0 for *p*_0_ = 3.4 and *p*_0_ = 4.2, respectively.

For a dynamic model like active spring-edge model, the response is dominated by a transient behavior. Figure D2(a) shows stress and strain as a function of time for this model without freezing the rest length degrees of freedom. As expected, the stress signal shows clear drifting behavior indicative of a transient state. Therefore, the Lissajous-Bowditch curve in Fig. D2(b) does not display an elliptic shape.

Due to the transient nature of dynamical models, we freeze their internal degrees of freedom— the rest lengths in this active spring-edge model—to calculate the storage and loss moduli (G′ and *G*″) at a minimum energy state. Consequently, we incorporate a *waiting time* (t_*w*_) parameter, representing the time allowed for the evolution of these models before their internal dynamics are halted.

**FIG. D2.**
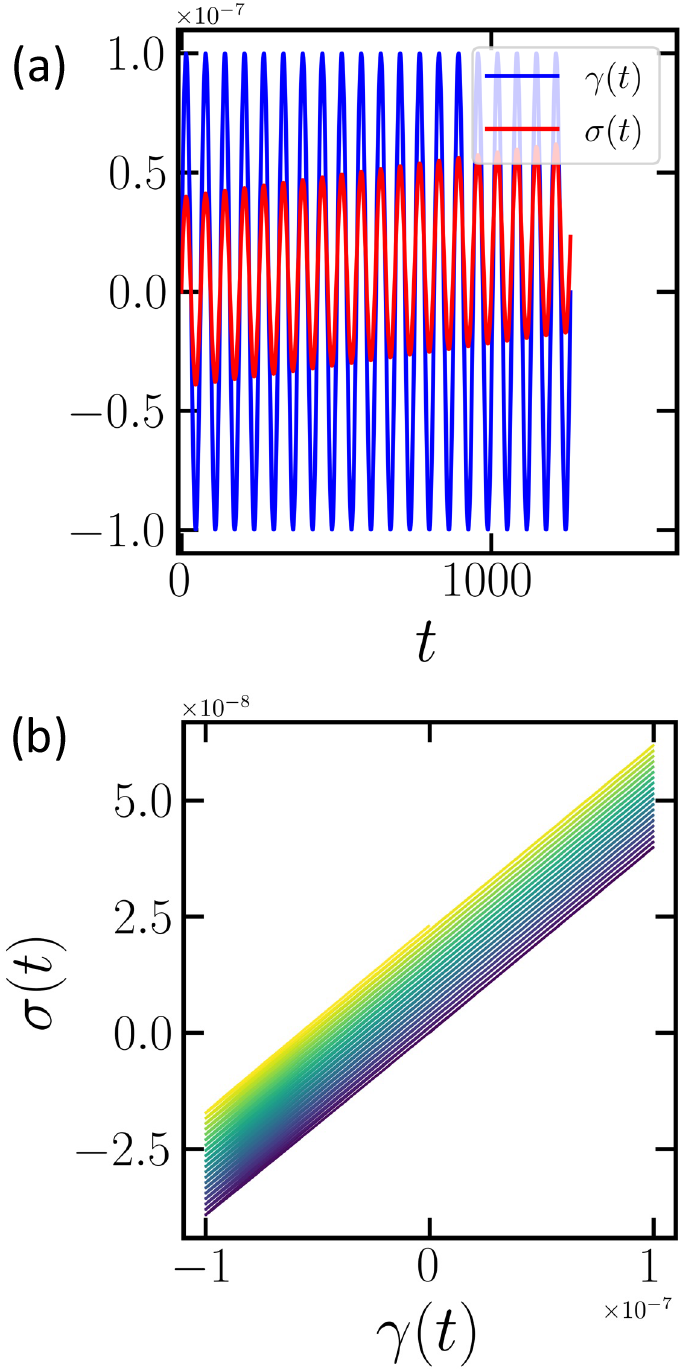
(a) Stress (red) and strain (blue) as a function of time for the active spring-edge model, where strain oscillations are applied without freezing the rest length degrees of freedom. Thus, there is a clear transient behavior in such dynamical model. (b) Lissajous-Bowditch curve derived from the data in (a) for the active spring-edge model, with lighter colors indicating later times.

## Appendix E: Rheological models

Here, we reproduce standard results on the rheology of spring-dashpot models for comparison to vertex model rheology simulations [16].

### 1. Zener/Standard Linear Solid Model

The Standard Linear Solid model, also known as the Zener model, provides a phenomenological representation of viscoelastic solids using a combination of spring and dashpot elements. This model, depicted in Fig. E1, consists of a spring arranged in parallel with a Maxwell element, which itself comprises a spring and a dashpot in series. Stress and strain in this system are denoted by *σ* and *ε* respectively, with the relationships *σ* = *kε* for springs and 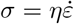 for dashpots.

For parallel components, the total stress is the sum of individual stresses (*σ*_total_ = *σ*_1_ + *σ*_2_) and the strain is equal across all components (*ε*_total_ = *ε*_1_ = *ε*_2_). For series components, the total strain is the sum of individual strains (*ε*_total_ = *ε*_1_ + *ε*_2_), and the stress is equal across components (*σ*_total_ = *σ*_1_ = *σ*_2_).

By applying these fundamental relationships, along with their time derivatives and the basic stress-strain relationships for the springs and dashpots, we can effectively model and analyze the behavior of viscoelastic materials [52].

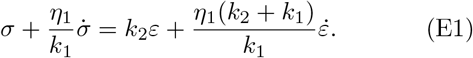

To derive the expression for the complex modulus *G*^*^ for the model given by the above equation, we start by assuming dynamic oscillations for strain and stress, where *ε*(t) = *ε*_0_*e*^*iωt*^ and *σ*(t) = *σ*_0_*e*^*iωt*^. Here, *ε*_0_ and *σ*_0_ are the amplitudes of strain and stress, respectively, and *ω* is the angular frequency. Substituting the expressions for *ϵ*(t) and *σ*(t) into the differential equation

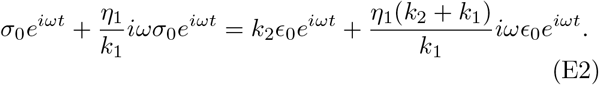

By solving for 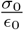 we find *G*^*^

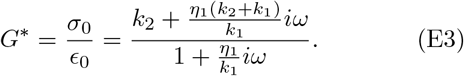

This expression provides the complex modulus *G*^*^ which encompasses both the storage modulus *G*′ and the loss modulus *G*″ based on its real and imaginary parts respectively. These are

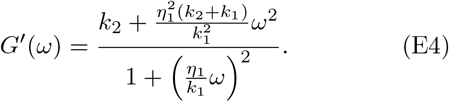

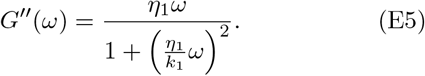

From these expressions, we derive the scaling behavior of *G*′ and *G*″ as the frequency *ω* approaches zero. In this limit, *G*′ reaches a constant plateau, while *G*″ ∼ *ω*.

### 2. Burgers/Standard Linear Liquid Model

The Standard Linear Liquid model, also known as Burgers model, provides a phenomenological representation of viscoelastic liquids using a combination of spring and dashpot elements. Using a similar approach as above, we can derive the following relation between stress and strain for this model

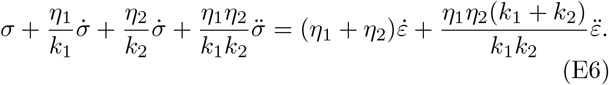

**FIG. E1.**
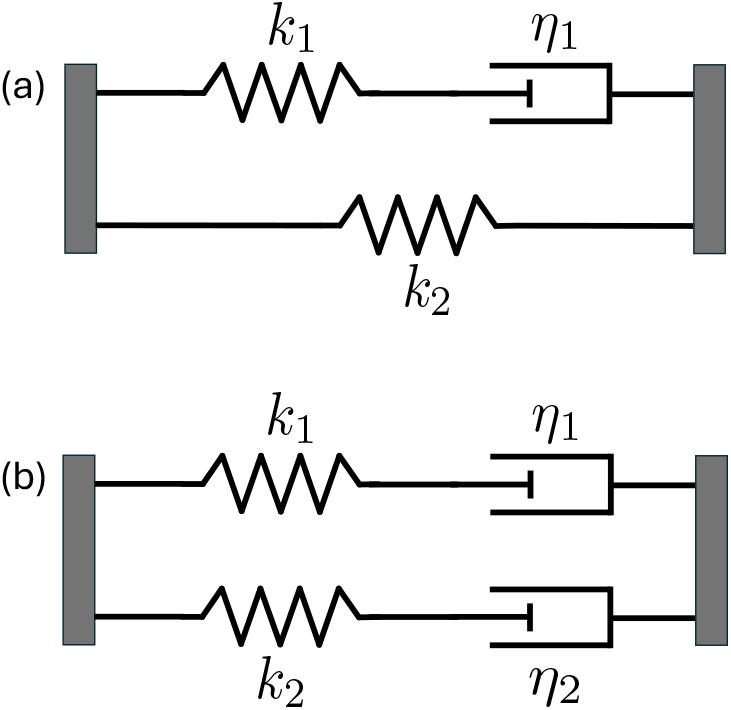
(a) Standard linear solid or Zener model. (b) Standard linear liquid or Burgers model

By substituting *ε* = *ε*_0_*e*^*iωt*^ and *σ*(t) = *σ*_0_*e*^*iωt*^ in the above equation, we can derive the expression for the complex modulus *G*^*^ = *σ*_0_*/ε*_0_ as

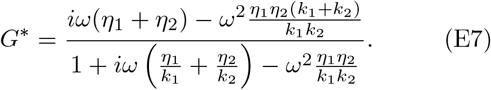

From this expression, we find *G*′ and *G*″ as

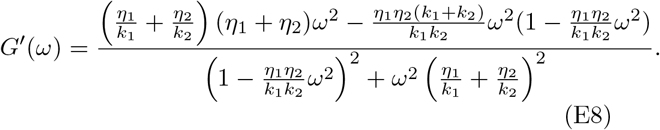

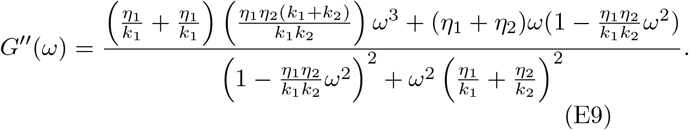

Therefore,in the low frequency limit *ω* → 0, this model exhibits *G* ∼ *ω*^2^ and *G*″ ∼ *ω*.

## Notes

### Competing Interest Statement

The authors have declared no competing interest.

### Summary of Updates

The manuscript was significantly edited to include new simulations and analysis demonstrating universality in the finite-frequency mechanical response of models for tissues, i.e. the viscoelastic behavior. This augments the original analysis in the manuscript focused on the zero-frequency response, i.e. the shear modulus.

